# SingleCellGGM enables gene expression program identification from single-cell transcriptomes and facilitates universal cell label transfer

**DOI:** 10.1101/2023.02.05.526424

**Authors:** Yupu Xu, Yuzhou Wang, Shisong Ma

## Abstract

Gene co-expression analysis of single-cell transcriptomes that aims to define functional relationships between genes is challenging due to excessive dropout values. Here, we developed a single-cell graphical Gaussian model (SingleCellGGM) algorithm to conduct single-cell gene co-expression network analysis. When applied to mouse single-cell datasets, SingleCellGGM constructed networks from which gene co-expression modules with highly significant functional enrichment were identified. We considered the modules to be gene expression programs (GEPs). These GEPs enable direct cell-type annotation of individual cells without cell clustering, and they are enriched with genes required for the functions of the corresponding cells, sometimes at a level greater than 10-fold. The GEPs are conserved across datasets and enable universal cell-type label transfer across different studies. We also proposed a dimension-reduction method through averaging-by-GEPs for single-cell analysis, enhancing the interpretability of results. Thus, SingleCellGGM offers a unique GEP-based perspective to analyze single-cell transcriptomes and reveals biological insights shared by different single-cell datasets.

## Introduction

Single-cell RNA-sequencing (scRNA-seq) technologies are now widely used, with more than 1,300 tools having been developed to conduct cell-based and gene-level analyses on single-cell datasets (Zappia et al. 2018). Cell-based analysis, which aims to identify cell types and trajectories of cellular development, has greatly enhanced our understanding of cellular heterogeneities (Trapnell et al. 2014; La Manno et al. 2018; Hao et al. 2021). However, gene-level analysis, which aims to define functional relationships between genes and identify gene regulators of cellular functions, remains challenging, and currently available tools can still be improved upon (Crow and Gillis 2018). Yet, such gene-level analysis is critical to unlocking the molecular logic of life.

Gene co-expression network (GCN) analysis is an effective method for identification of functional relationships between genes. However, such analysis is challenging when dealing with single-cell transcriptome datasets. In GCN analysis, Pearson correlation coefficients (*r*) are calculated between genes based upon their expression to determine whether they are co-expressed. According to the “guilt by association” paradigm, co-expressed genes tend to function within the same pathways. GCN analysis has been successfully applied to bulk transcriptome datasets to identify gene co-expression modules functioning in diverse pathways (Stuart et al. 2003; Langfelder and Horvath 2008; Mentzen and Wurtele 2008). The identified modules can be considered as gene expression programs (GEPs). However, similar analysis of single-cell transcriptomes is challenging because co-expression signals are often strikingly weak in single-cell datasets (Crow and Gillis 2018). Single-cell datasets contain frequent dropout expression values, which obscures the calculation of *r* and makes most gene-level analyses a challenge. One method of addressing dropouts is grouping cells with similar expression to form metacells and then performing co-expression analysis, as proposed by MetaCell and scCorr (Baran et al. 2019; Xu et al. 2022a). Methods including MAGIC and scImpute have also been developed to impute expression values of dropouts to improve downstream gene-level analysis (Li and Li 2018; van Dijk et al. 2018). In addition to *r*, the partial correlation coefficient (*pcor*) is also adopted by methods such as SILGGM and scLink in co-expression analysis of single-cell datasets (Zhang et al. 2018; Li and Li 2021). *Pcor* is the direct correlation between two genes after removing the effects of other genes, which is considered a more representative coefficient than *r* for gene network analysis (Wille et al. 2004; Schafer and Strimmer 2005). The co-expression network based on *pcors* is known as the graphical Gaussian model (GGM). However, these methods have only been tested in small-scale datasets, and have not been widely adopted by the single-cell research community.

Non-negative matrix factorization (NMF) is another method for identification of GEPs from single-cell datasets. NMF was originally developed to extract features from computer graphics, but it was quickly adopted for microarray data analysis and more recently for single-cell analysis (Lee and Seung 1999; Brunet et al. 2004; Kotliar et al. 2019; Elyanow et al. 2020). For a single-cell gene expression matrix, termed A, NMF can factorize it into two non-negative matrices termed W and H, with W specifying linear combinations of genes that can be treated as GEPs and H specifying the GEPs’ usage levels within individual cells. Kotliar et al. developed a consensus NMF (cNMF) method for analysis of two single-cell datasets from brain organoid and visual cortex samples, and they identified two types of GEPs: identity GEPs with cell-type specific usages, and activity GEPs with shared usages across different cell types which represent common pathways among them (Kotliar et al. 2019). cNMF has recently been applied to identify GEPs functioning in rove beetle defense glands and during mouse spermatogenesis (Bruckner et al. 2021; Xu et al. 2022b).

Here, we have developed a single-cell graphical Gaussian model (SingleCellGGM) algorithm to conduct single-cell gene co-expression network analysis. When applied to two single-cell datasets from mice, SingleCellGGM generated co-expression networks from which gene co-expression modules with significant functional enrichment were identified, and we considered these modules as GEPs. The GEPs mostly have cell-type specific expression and allow for direct cell-type annotation of individual cells without cell clustering. They are also enriched with genes required for the functioning of the corresponding cell types. The GEPs are conserved across datasets and enable universal cell type label transfer across different studies. We also proposed a dimension reduction method via averaging-by-GEPs for single-cell analysis, allowing for enhanced interpretability of analysis results. SingleCellGGM thus provides a unique GEP-based solution to analyze single-cell transcriptomes.

## Results

### An algorithm for single-cell gene co-expression network analysis

We developed an algorithm called single-cell graphical Gaussian model (SingleCellGGM) to conduct single-cell gene co-expression network analysis (**Figure 1A**). SingleCellGGM is modified from a previous method for GGM gene co-expression network analysis of bulk transcriptomes (Ma et al. 2007; Wang et al. 2023). It uses a process consisting of ∼20,000 iterations to calculate partial correlation coefficients (*pcors*) between genes. In each iteration, 2,000 genes are randomly chosen and used for *pcor* calculation. Throughout the process, each gene pair is chosen in multiple iterations with multiple *pcors* calculated, and the *pcor* with the lowest absolute value is selected as the final *pcor* for that gene pair. SingleCellGGM then retains those *pcors* ≥ a selected cutoff value for downstream analysis. As single-cell datasets are relatively sparse with many dropout values, SingleCellGGM further removes gene pairs that are co-expressed in less than a selected number of (e.g., 10) cells. Thus, SingleCellGGM uses all gene pairs with *pcors* ≥ a cutoff value and co-expressed in ≥ a cutoff number of cells to build a final gene co-expression network.

**Figure 1.**
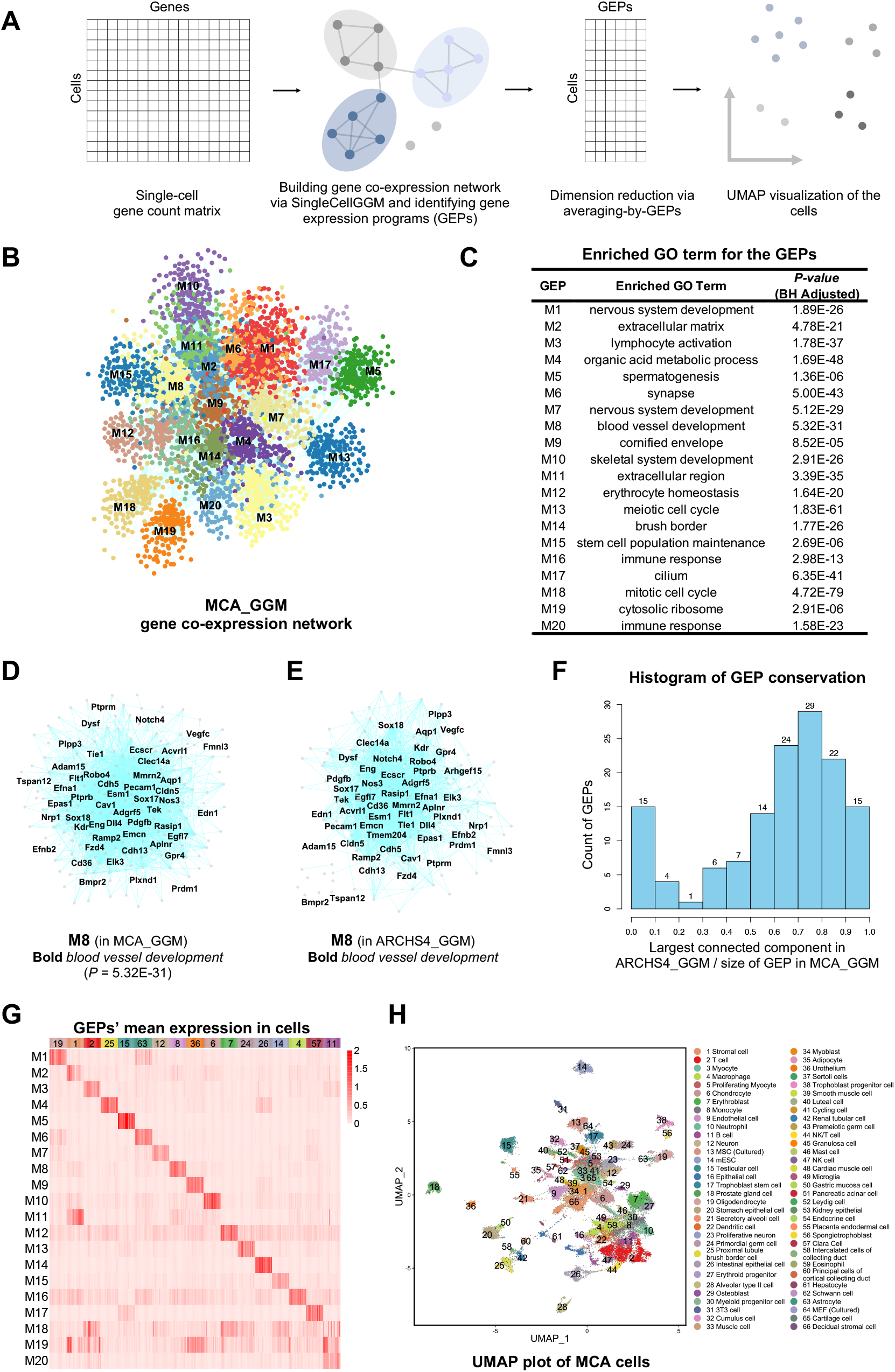
Single-cell gene co-expression network analysis using SingleCellGGM identifies gene expression programs from the MCA dataset. **(A)** The analysis workflow. **(B)** MCA_GGM gene co-expression network. Nodes represent genes, and connections between nodes indicate co-expression. Nodes are colored according to their gene expression program (GEP) identities. Only genes from the first 20 largest GEPs are shown due to space limitations. **(C)** Enriched GO terms for the GEPs. Only the first 20 GEPs are shown due to space limitations. **(D)** A subnetwork for GEP M8 of MCA_GGM. The subnetwork contains 179 genes. Gene names are shown for a portion of genes due to space limitations. In this and other subnetworks, genes possessing specific Gene Ontology (GO) or Mammalian Phenotype Ontology (MP) terms are highlighted in bold, italic, or red. *P*, enrichment *P* value. **(E)** A subnetwork extracted from ARCHS4_GGM for the genes contained within GEP M8 of MCA_GGM. The largest connected component of the subnetwork contains 157 genes, indicating that among the 179 genes in GEP M8 of MCA_GGM, 87.7% form a co-expression module in ARCHS4_GGM. **(F)** A histogram illustrating the percentage of genes within the MCA GEPs that also form modules within ARCHS4_GGM. **(G)** An expression heatmap for the first 20 GEPs (rows) in the cells sampled from different cell types (columns) in MCA. The cell type identities are indicated by the same number and color scheme as in (H). **(H)** A UMAP plot of MCA cells after dimension reduction via averaging-by-GEPs. The average expression values of GEPs in each cell were used as inputs for UMAP analysis. The cells are colored according to their cell type labels as annotated by the original MCA study.

We tested our SingleCellGGM algorithm on a mouse cell atlas (MCA) dataset (Han et al. 2018). The MCA dataset used a Microwell-seq platform to profile mouse single-cell transcriptomes. This dataset contains expression values of 25,133 genes in 61,637 cells sampled from across 43 tissues, as described in Figure 2 of the original publication. To enable comparisons with other datasets, we chose 24,802 genes with Ensembl gene IDs for network construction. After SingleCellGGM analysis, we obtained 127,229 gene pairs with *pcors* ≥0.03 and co-expressed in ≥10 cells (**Figure S1, Table S1**). These gene pairs were used to construct a GGM gene co-expression network named MCA_GGM. The network was clustered using the MCL algorithm (Enright et al. 2002), and 137 gene co-expression modules containing ≥10 genes were identified. As the genes within modules were co-expressed and may be regulated by the same transcriptional regulators, we considered these gene co-expression modules as gene expression programs (GEPs) (**Figure 1B, Table S2**). Gene Ontology (GO) analysis revealed that 123 of these GEPs had enriched GO terms (*P* ≤ 0.05, Benjamini‒Hochberg (BH) adjusted; **Figure 1C**, see **Table S3** for a complete list). For example, GEP M1 was enriched with 120 genes containing the GO term *nervous system development* (*P* = 1.89E-26), M2 with 38 genes for *extracellular matrix* (*P* = 4.78E-21), and M3 with 61 genes for *lymphocyte activation* (*P* = 1.78E-37). The results indicated that GEPs effectively grouped genes participating in the same biological processes or those from the same cellular compartments.

**Figure 2.**
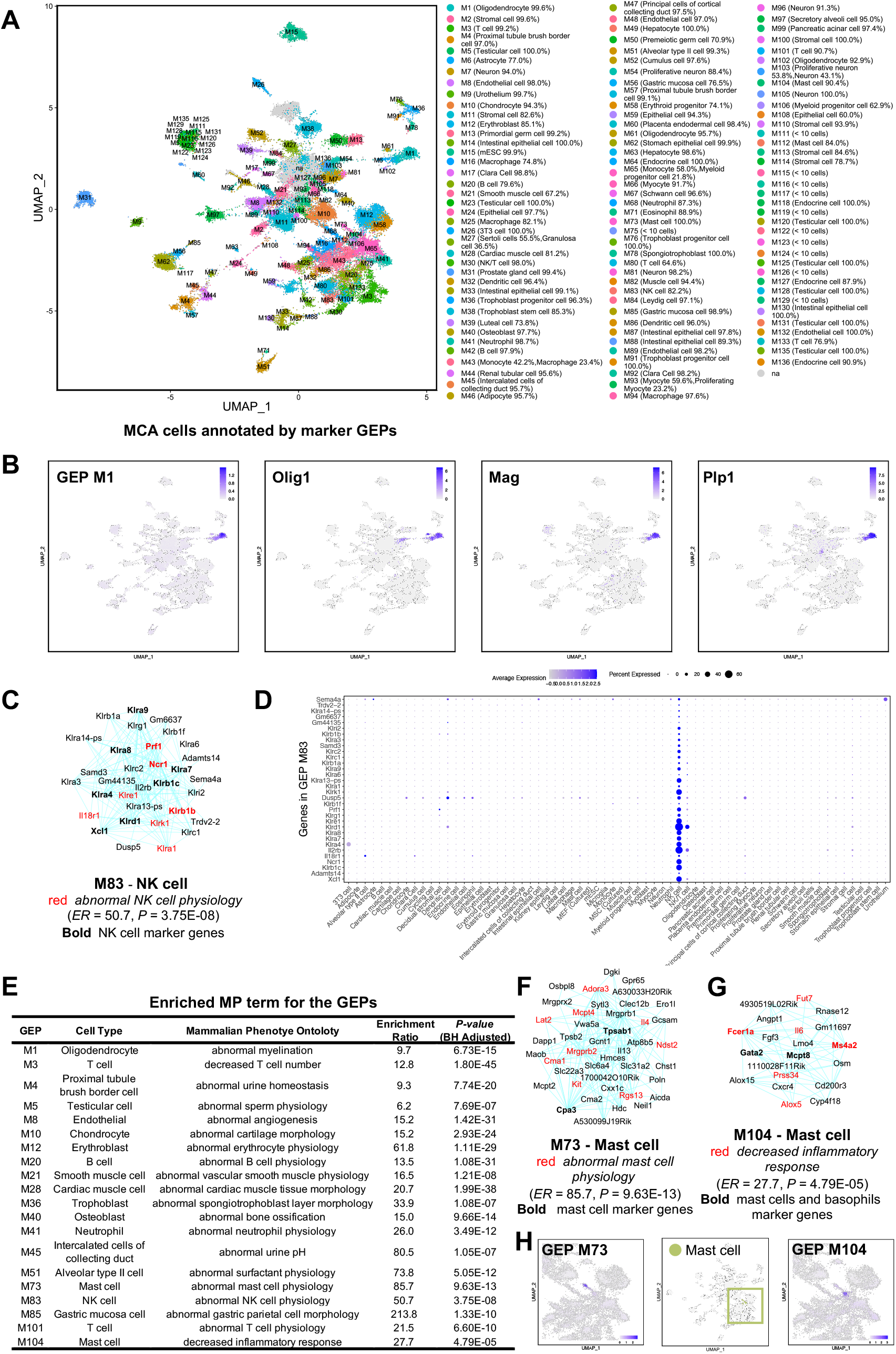
The identity GEPs identified from MCA_GGM. **(A)** The MCA cells annotated according to their marker GEPs. Among the identity GEPs expressed in a cell, the one with the highest average expression value was considered to be the marker GEP for the cell. A neighboring voting procedure was used to propagate marker GEP labels to neighboring unannotated cells. Cells are colored according to their marker GEPs. Cell type compositions of cells annotated by each GEP are shown in brackets. **(B)** Expression of GEP M1 is similar to that of *Olig1, Mag*, and *Plp1* in MCA cells. The expression values of GEPs are defined as their genes’ average expression values. *Olig1, Mag*, and *Plp1* are known oligodendrocyte marker genes contained within GEP M1. The color scale for each plot represents normalized and log-transformed expression values for the indicated GEP or genes, same as blow. **(C)** A subnetwork for GEP M83. The genes possessing the MP term *abnormal NK cell physiology* are enriched 50.7-fold within this GEP. *ER*, enrichment ratio. **(D)** Cell-type-specific expression of the genes in GEP M83. **(E)** The MP terms enriched within the GEPs. The results are shown for a portion of the GEPs. Refer to Table S6 for a full list of the results. **(F and G)** Two subnetworks for M73 and M104, marker GEPs for mast cells. **(H)** Expression of GEP M73 and M104 in MCA cells. Mast cells are highlighted on the map near the middle, and two enlarged areas containing mast cells depict the expression of M73 and M104.

The identified GEPs are robust and their genes form co-expression modules across other single-cell datasets or bulk RNA-seq samples. We constructed another two GGM co-expression networks, one using the ARCHS4 (v10) dataset with 91,658 mouse bulk RNA-seq samples, and the other using the Tabula Muris Senis (TMS) FACS single-cell dataset with 110,824 cells (Lachmann et al. 2018; Tabula Muris 2020), and named them ARCHS4_GGM and TMS_GGM (**Table S4**). The genes from MCA GEPs form modules in ARCHS4_GGM and TMS_GGM. For example, GEP M8 contains 179 genes and functions primarily in *blood vessel development* (*P* = 5.32E-31; **Figure 1D**). We extracted two subnetworks for these 179 genes from ARCHS4_GGM and TMS_GGM and found that their largest connected components consisted of 157 and 147 genes, respectively (**Figure 1E and Figure S2A**). Therefore, 87% and 82% of genes in M8 form gene modules in ARCHS4_GGM and TMS_GGM, respectively. Among 137 GEPs from MCA_GGM, 92 have at least 60% of their genes also form modules in ARCHS4_GGM and/or TMS_GGM, indicating that these GEPs are highly conserved in bulk RNA-seq samples or the TMS dataset, providing supporting evidence for their biological significance (**Figure 1F, Figure S2B and Table S4**). Among the 13 GEPs with less than 10% genes which form conserved modules, 12 of them (M111, M115, M116, M119, M122, M123, M125, M126, M128, M129, M131, and M135) have specific expression in testicular cells (see below for expression analysis of the GEPs).

### A dimension-reduction method based on averaging by GEPs

We examined the expression patterns of the genes within the MCA GEPs. Based upon cell types annotated by the original MCA study (Han et al. 2018), we uncovered that a large portion of the GEPs have cell-type-specific expression patterns. Individual genes from these GEPs show sparse but clear, specific expression in certain cell types, for example, GEP M1 in oligodendrocytes, M2 in stromal cells, and M3 in T cells (**Figure S3**). GEPs with ubiquitous expression across different cell types are also present, such as M18 and M19. These two GEPs have enriched GO terms for *mitotic cell cycle* (*P* = 4.72E-79) and *cytosolic ribosome* (*P* = 2.91E-06) respectively, indicating that they function in housekeeping processes including cell cycle regulation and protein biosynthesis. Expression patterns observed in individual genes from these GEPs are sparse and mosaic, which may be due to frequent dropout values in single-cell datasets. When using the average expression values of each GEP within every cell to draw a heatmap, cell-type-specific expression patterns within the GEPs became clearer (**Figure 1G**).

Considering our above observation, we wished to determine whether gene expression values of the cells within the MCA could be summarized as average gene expression values within each of the 137 GEPs. Such summarization is analogous to the dimension reduction technique like principal component analysis (PCA), where the expression of thousands of genes in cells is summarized by a small number of principal components (PCs) (Hao et al. 2021). The PC embeddings of cells are further processed by the uniform manifold approximation and projection (UMAP) or t-distributed stochastic neighbor embedding (tSNE) dimensionality reduction methods to generate a map for cell visualization and clustering (van der Maaten and Hinton 2008; Becht et al. 2018). However, as PCs are linear combinations of all genes, it can be difficult to extract biological meanings from them. Here, we calculated the average expression values of genes within each of the 137 GEPs for every cell. A UMAP analysis using these average expression values as inputs was performed to generate a map which clearly separated the different cell types in the MCA dataset (**Figure 1H**). The map also preserves higher order structure of the cell types: the cells from the hematopoietic lineage, neuro system, stem cell and reproductive system, and those related to epithelia tend to form large groups within the map. Therefore, averaging-by-GEPs can be used as a dimensionality reduction technique. Compared to PCA, an advantage of the GEP-based reduction approach is its interpretability, with the ability to assign biological functions to the GEPs, as demonstrated below.

The above analysis indicated that GEPs identified from MCA have functional significance, as exemplified by their enriched GO terms, their conservation in bulk transcriptomes and other single-cell datasets, and their cell-type specific expression patterns. Based on their expression patterns and functional annotations, we divided these GEPs into three categories: 1) 114 identity GEPs, which have specific expression in one or a limited number of cell types; 2) 18 activity GEPs, which mostly have ubiquitous expression and shared functions across many cell types; 3) Five morphogenesis GEPs, which are expressed chiefly in fetal or neonatal tissues and function in regulating cell differentiation and body morphogenesis (**Table S5**).

### Identity GEPs enable direct annotation of individual mouse cells

The 114 identity GEPs can be used to annotate individual cells for their cell types. In a typical single-cell analysis, cells are first divided into clusters, and the clusters are then annotated for their cell types depending upon the expression of marker genes. We uncovered that the identity GEPs can be used as marker GEPs for annotation of individual cells directly. For every cell in the MCA, we determined which identity GEPs are expressed in that cell. Briefly, if a GEP has an average expression value in a cell ≥1/2 of the GEP’s maximal average expression value among all cells, we considered the GEP to be expressed in that cell. Among all identity GEPs expressed in a cell, the one with the highest average expression value was considered to be the marker GEP for the cell. As only a portion of cells were assigned marker GEPs in this initial step, we used a neighbor voting procedure to assign GEPs to the cells without marker GEPs but connected to multiple cells with the same marker GEPs. The cells were annotated according to their marker GEPs. We discovered that the cell marker GEP annotations largely overlapped with their original MCA cell-type labeling, and most marker GEPs had cell-type specificities (**Figure 2A and Table S5**). For example, among the 1,125 cells annotated by GEP M1, 1,120 (99.6%) are oligodendrocytes; 99.6% of the 805 cells annotated by M2 are stromal cells; 99.2% of the 1,886 cells annotated by M3 are T cells; and 97% of the 733 cells annotated by M4 are proximal tubule brush border cells. Therefore, we can assign cell-type annotations to individual cells according to their marker GEPs. This annotation relies on the average expression values of multiple genes within the GEPs and can effectively mitigate the influence of dropout expression values.

Additionally, most marker GEPs contain multiple known marker genes of the corresponding cell types (**Table S5**), and the expression of the marker GEPs, which are defined as the average expression values of all the genes within the GEPs, often matches the expression of the marker genes. As an example, M1, a marker GEP for oligodendrocytes, contains the known oligodendrocyte marker genes *Olig1, Mag*, and *Plp1*. The expression pattern of GEP M1 is similar to those of *Olig1, Mag*, and *Plp1* (**Figure 2B**). This observed similar expression between the GEPs and known marker genes provides further support that GEPs can be used as markers for cell annotation.

The marker GEPs are also enriched with genes required for the functions of the corresponding cells. For example, M83 is a marker GEP for natural killer (NK) cells, containing 32 genes that are mainly expressed in NK cells, including 10 known NK cell marker genes (**Figure 2C, D**). According to the Mouse Genome Information (MGI) database, 7 of these 32 genes possess the Mammalian Phenotype Ontology (MP) term *abnormal NK cell physiology*, for example, *Ncr1* and *Klra1* (**Figure 2C**) (Narni-Mancinelli et al. 2012; Eppig et al. 2014; Bern et al. 2017). As MP term annotations are assigned to genes based on the phenotypes of their mouse mutants, the 7 genes with the MP term *abnormal NK cell physiology* have NK cell-related functions. Throughout all 24,802 genes used for network construction, only 107 genes possess the same MP term; the term is thus enriched 50.7-fold within GEP M83 (*Enrichment Ratio, ER* = 50.7, *P* = 3.75E-08; see **Table S6** for MP enrichment analysis results for all GEPs). As another two examples, M40, a marker GEP for osteoblasts, is enriched with 18 genes for *abnormal bone ossification* (*ER* = 15.0, *P* = 9.66E-14), and M51, a marker GEP for alveolar epithelial type II (AT2) cells, is enriched with 9 genes for *abnormal surfactant physiology* (*ER* = 73.8, *P* = 5.05E-12) (**Figure 2E**). Since NK cell-, osteoblast-, and AT2 cell-related genes are highly enriched within these three GEPs, according to the “guilt by association” paradigm, we speculate that other genes within these GEPs may also function across similar pathways and are therefore ideal candidates for future investigations. Correspondingly, marker GEPs and candidate genes were found for other cells, such as chondrocytes (M10), erythroblasts (M12), neutrophils (M41), and intercalated cells of collecting duct (M45), which will facilitate functional studies of these cells.

### Identity GEPs reveal different states or subtypes of the same cell types

The identity GEPs also reveal different cellular states or subtypes of the same cell types. For example, we identified two marker GEPs for mast cells, M73 and M104, which are expressed in two different mast cell populations (**Figure 2F-H**). We speculate that the cells annotated by M73 are mature mast cells, while those annotated by M104 are mast cell progenitors. M73 contains mature mast cell marker genes *Cpa3* and *Tpsab1*, which encode carboxypeptidase A3 and tryptase alpha-1/beta-1 that function as mast cell mediators (Moon et al. 2014). Conversely, M104 contains *Fcer1a, Ms4a2*, and *Mcpt8*, which are marker genes for both mast cells and basophils. Among them, *Mcpt8* encodes a protease expressed in the early stage of mast cell development (Hamey et al. 2021). This GEP also contains *Gata2*, encoding a transcription factor (TF) required for differentiation and maintenance of mast cells and basophils (Li et al. 2015). Considering that a portion of cells annotated by M104 originated from bone marrow (**Figure S4**), these cells may be common progenitors for both mast cells and basophils. Additionally, M73 and M104 contained 9 and 6 genes with the MP terms *abnormal mast cell physiology* and *decreased inflammatory response*, representing an 85.7- and 27.7-fold enrichment over the background, respectively (*P* = 9.63E-13 and 4.79E-05). Therefore, these two GEPs’ uncharacterized genes are also ideal candidate genes for studying mast cell functions.

Multiple GEPs were also identified related to other cells, such as B cells (M20, M42), dendritic cells (M32, M86), neurons (M7, M81, M96, M103, M105), oligodendrocytes (M1, M6, M61, M102), stromal cells (M2, M11, M100, M110), and gastric mucosa cells (M56, M85). These GEPs represent different states or subtypes of the cells, and they identify genes that differentiate these states or subtypes, providing an ample number of candidates to study underlying molecular mechanisms.

### Activity GEPs identify shared transcription programs across cell types

In addition to GEPs with cell type specificities, SingleCellGGM network analysis was also able to identify 15 GEPs with shared expression across many different cell types, which are considered activity GEPs (**Table S5**). These GEPs regulate the cell cycle and DNA replication (M18, M34, M121), responses to virus and stress (M35, M53, M74), RNA processing (M134), protein biosynthesis (M19, M37), mitochondrial activities (M79), endoplasmic reticulum-related activities (M90, M107), cytoskeleton organization (M69), cholesterol biosynthesis (M109), and some with unknown functions (M29). Three other GEPs, also regulating mitochondrial activities (M22 and M55) and RNA processing (M98), are considered activity GEPs based on their functional annotations.

Among these, M18 is a GEP regulating mitotic cell cycle activity, enriched with 82 genes for *mitotic cell cycle* (*P* = 4.72E-79), such as *Aurka, Ccna1*, and *Cdk1*. The GEP is expressed across many different cell types (**Figure 3A, B**). Similarly, M34 is a GEP modulating *DNA replication* (*P* = 4.50E-48), also with expression across a wide range of cell types (**Figure 3C, D**). These two GEPs are enriched with genes possessing the MP terms *abnormal mitosis* and *chromosomal instability*, respectively (*ER* = 28.6 and 22.5, *P* = 1.16E-18 and 4.14E-07, **Table S6**). We used GEP M18 to evaluate cell cycle activity. We considered a cell to be in an active cell cycle if its GEP M18 average expression value was ≥1/2 of the GEP’s maximal average expression value across all cells and calculated a cell cycle index for each cell type reflecting what fraction of its cells were in an active cell cycle. We also used GEP M34 to calculate a DNA replication index. Interestingly, we discovered that some cell types had different rankings of the two indices; for example, erythroblasts and erythroid progenitors had the 2 highest DNA replication indices, while their cell cycle indices ranked only 14th and 22nd among all cell types (**Figure 3E, F**). This indicates that DNA replication may be decoupled from cell cycle regulation in these two cell types. In addition to cell cycle and DNA replication GEPs, other activity GEPs may also be used for evaluation of cell status. These include RNA processing activity (M98, M134), protein biosynthesis (M19, M37), mitochondrial activities (M22, M55, M79), ER-related activities (M90, M107), cytoskeleton-related activities (M69), and cholesterol biosynthesis rate (M109).

**Figure 3.**
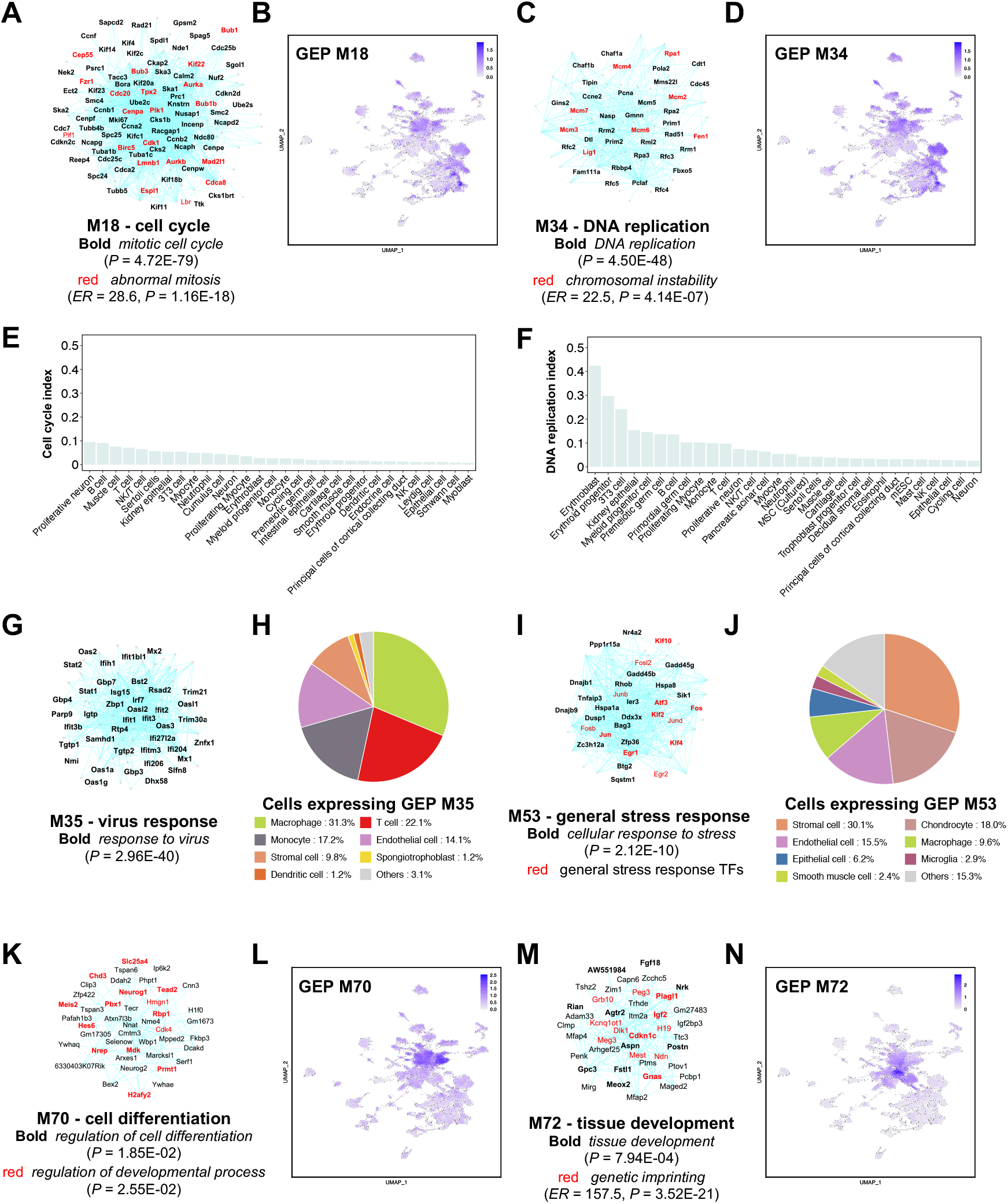
The activity and morphogenesis GEPs identified from MCA_GGM. **(A-D)** Two subnetworks of activity GEPs M18 and M34 and their expression in MCA cells. **(E and F)** Cell cycle index and DNA replication index for cell types in MCA. Only the top 30 cell types are shown. **(G - J)** Two subnetworks of activity GEPs M35 and M53 and the cells expressing these two GEPs. **(K - N)** Two subnetworks of morphogenesis GEPs M70 and M72 and their expression in MCA cells.

We also identified an activity GEP, M35, involved in the immune response to viruses. This GEP is enriched with 40 genes for *response to virus* (*P* = 2.96E-40), including 12 interferon-induced genes and 3 TF genes, *Stat1, Stat2*, and *Irf7*, all of which are critical for virus response (Mogensen 2018) (**Figure 3G**). Interestingly, the GEP is expressed across several cell types, including macrophages, T cells, monocytes, endothelial cells, and stromal cells (**Figure 3H and Figure S5A**). Another GEP, M53, is considered to function in general stress response, as it is enriched with genes for *cellular response to stress* (*P* = 2.12E-10) and contains key TFs which function in stress responses, such as *Fos, Fosb, Jun, Junb, Jund, Egr1*, and *Atf3* (Denisenko et al. 2020) (**Figure 3I**). This GEP is ubiquitously expressed in stromal cells, chondrocytes, endothelial cells, macrophages, epithelial cells, microglia, and smooth muscle cells (**Figure 3J and Figure S5B**). We envision that these two GEPs can be used to monitor cell status on virus infection and stress loading.

### Morphogenesis GEPs reveal programs regulating cell differentiation and tissue patterning

In addition to identity and activity GEPs, we also identified a third category of GEPs that are highly expressed in fetal or neonatal tissues. These GEPs have enriched functions in regulation of cell differentiation and tissue patterning, and we therefore considered them as morphogenesis GEPs. For example, M70 is a GEP enriched with 12 genes for *regulation of cell differentiation* and 14 genes for *regulation of developmental process* (*P* = 1.85E-02 and 2.55E-02), such as *Meis2, Neurog1*, and *Tead2* (**Figure 3K**). It is highly expressed in fetal or neonatal tissues, such as the embryonic mesenchyme, fetal brain, fetal kidney, fetal intestine, fetal lung, and neonatal brain (**Figure S6A**). In terms of cells, the GEP is highly expressed in (proliferative) myocytes, (proliferative) neurons, epithelial cells, muscle cells, and cycling cells. (**Figure 3L and S6B**). As another example, M72 is a GEP enriched with 14 genes specifically for *tissue development* (*P* = 7.94E-04), such as *Agtr2, Fgf18*, and *Igf2* (**Figure 3M**). This GEP is highly expressed in neonatal muscle, neonatal skin, and neonatal rib, as well as in stromal cells, myocytes, and chondrocytes (**Figure 3N and S6C, D**). Therefore, it is clear that both GEPs do not have cell type specificity and instead may function in regulating cell differentiation and tissue patterning in the early stage of mouse development. Interestingly, M72 is also enriched with 12 genes containing the MP term *genetic imprinting* (*ER* = 157.5, *P* = 3.52E-21), which is consistent with the notion that epigenetic regulation is important for tissue development. Another three GEPs, M77, M95, and M137, have biased expression in (proliferative) myocytes and/or stromal cells of fetal kidney, fetal intestine, and fetal lung (**Figure S6E-M**). They are enriched with genes marked for *pattern specification process, embryonic morphogenesis*, and *embryonic skeletal system morphogenesis* (*P* = 3.24E-09, 1.46E-08, and 1.64E-05, respectively). Therefore, we also considered them to be morphogenesis GEPs.

### SingleCellGGM outperforms other tools in identification of GEPs

In addition to SingleCellGGM, other tools have also been developed to conduct gene co-expression network analysis, such as ppcor and SILGGM (Kim 2015; Zhang et al. 2018). ppcor and SILGGM also calculate *pcors* between genes to construct gene co-expression networks. Therefore, we used ppcor and SILGGM to construct two gene co-expression networks for MCA (**Table S7**) and clustered the networks via the MCL algorithm to obtain GEPs. We then compared SingleCellGGM with ppcor and SILGGM to evaluate their performance in identifying GEPs drawing upon the same dataset. While SingleCellGGM network analysis identified 137 GEPs, 123 of which had enriched GO terms, ppcor and SILGGM identified 224 and 258 GEPs, respectively, 98 and 175 of which had significantly enriched GO terms (*P* ≤ 0.05; **Table S8**). However, the sizes of the GEPs identified by ppcor and SILGGM are much smaller than those identified by SingleCellGGM. SingleCellGGM identified 64 significant GEPs with enriched GO terms and containing ≥50 genes, while ppcor and SILGGM only identified 1 and 5, respectively. The GEPs with enriched GO terms in SingleCellGGM contained 8,895 genes in total, while those in ppcor and SILGGM contained only 2,294 and 3,758 genes, respectively (**Figure 4**). With this in mind, SingleCellGGM could potentially provide functional annotations to more genes based on the identified GEPs than the other two methods. Additionally, if the *P* value cutoff was more stringent and set to 1E-20, SingleCellGGM, ppcor, and SILGGM would identify 32, 2, and 6 significant GEPs, respectively (**Table S8**). Taken together, SingleCellGGM performs more optimally than ppcor and SILGGM in identifying GEPs for gene function analysis.

**Figure 4.**
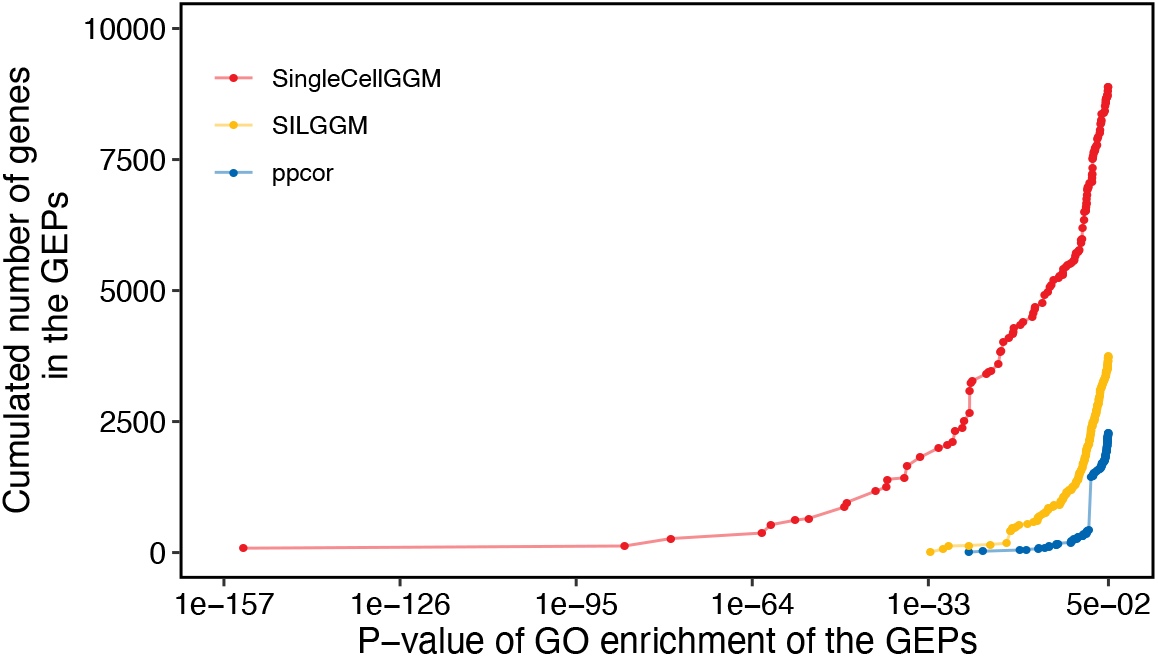
A comparison between SingleCellGGM, SILGGM, and ppcor. SingleCellGGM was compared to both SILGGM and ppcor in their ability to identify GEPs from MCA. The plot shows the cumulative number of genes within the GEPs with *P* values less than or equal to the indicated *P* values identified by SingleCellGGM, SILGGM, and ppcor. Each dot represents a GEP. At a given *P* value cutoff, the GEPs identified by SingleCellGGM contain more genes than those identified by SILGGM and ppcor.

We compared SingleCellGGM to the cNMF method in identifying GEPs from the MCA dataset (Kotliar et al. 2019). cNMF uses non-negative matrix factorization to identify GEPs from single-cell transcriptomes. Similar to SingleCellGGM, cNMF identified GEPs with cell-type specific usages from the MCA dataset. Indeed, almost all GEPs identified by cNMF were specific to one or a limited number of related cell types, and only 1 GEP displayed shared usages across many different cell types and thus can be considered as an activity GEP (**Figure S7**). This GEP contained marker genes (genes that are statistically associated with the GEP as determined by cNMF) functioning in *cellular response to stress* (*P* = 5.06E-08), such as *Atf3, Egr1, Fos*, and *Jun*, which is similar to GEP M53 from MCA_GGM. In contrast, among the 18 activity GEPs identified by SingleCellGGM, 15 demonstrated shared expression across many cell types, including those involved in cell cycle regulation, protein biosynthesis, and endoplasmic reticulum-related activities, however, similar shared GEPs were not identified by cNMF. Thus, cNMF does not operate ideally in identifying activity GEPs which are shared across different cell types from the MCA dataset, and SingleCellGGM performs more optimally in this regard.

### Analysis of a mouse brain single-cell dataset identifies GEPs for different neurons

We also used SingleCellGGM to analyze a mouse brain single-cell dataset obtained via the 10X Genomics platform (La Manno et al. 2021). The dataset contains 292,495 cells from developing mouse brains sampled daily between embryonic day (E)7 and E18. The original analysis divided the cells into 23 classes and 133 subclasses according to their cell types, with many subclasses corresponding to neurons from different brain regions. SingleCellGGM identified 73,786 gene pairs with *pcors* ≥0.03 and co-expressed in ≥10 cells from the dataset, and these were used to build a co-expression network named MouseBrain_GGM (**Table S9**). Using MCL clustering, 158 gene co-expression modules with ≥10 genes were identified in the network, and they were considered GEP B1 to B158. Among them, 135 had enriched GO terms (*P* ≤ 0.05; **Table S10, S11**). A UMAP analysis using average gene expression values from the GEPs in each cell as inputs differentiated the cells according to their original class and subclass labels (**Figure 5A and Figure S8**). The map depicts a continuum of cells from neural tubes to radial glia, neuroblasts, and neurons. The radial glia, neuroblasts and neurons from the forebrain, midbrain, and hindbrain are clustered according to their region of origins (**Figure 5B**). Based on their expression patterns, we divided the 158 GEPs into 116 identity GEPs, 25 activity GEPs, and 17 morphogenesis GEPs (**Table S12**). We focused on the identity GEPs to identify transcriptional signatures of different mouse brain cells.

**Figure 5.**
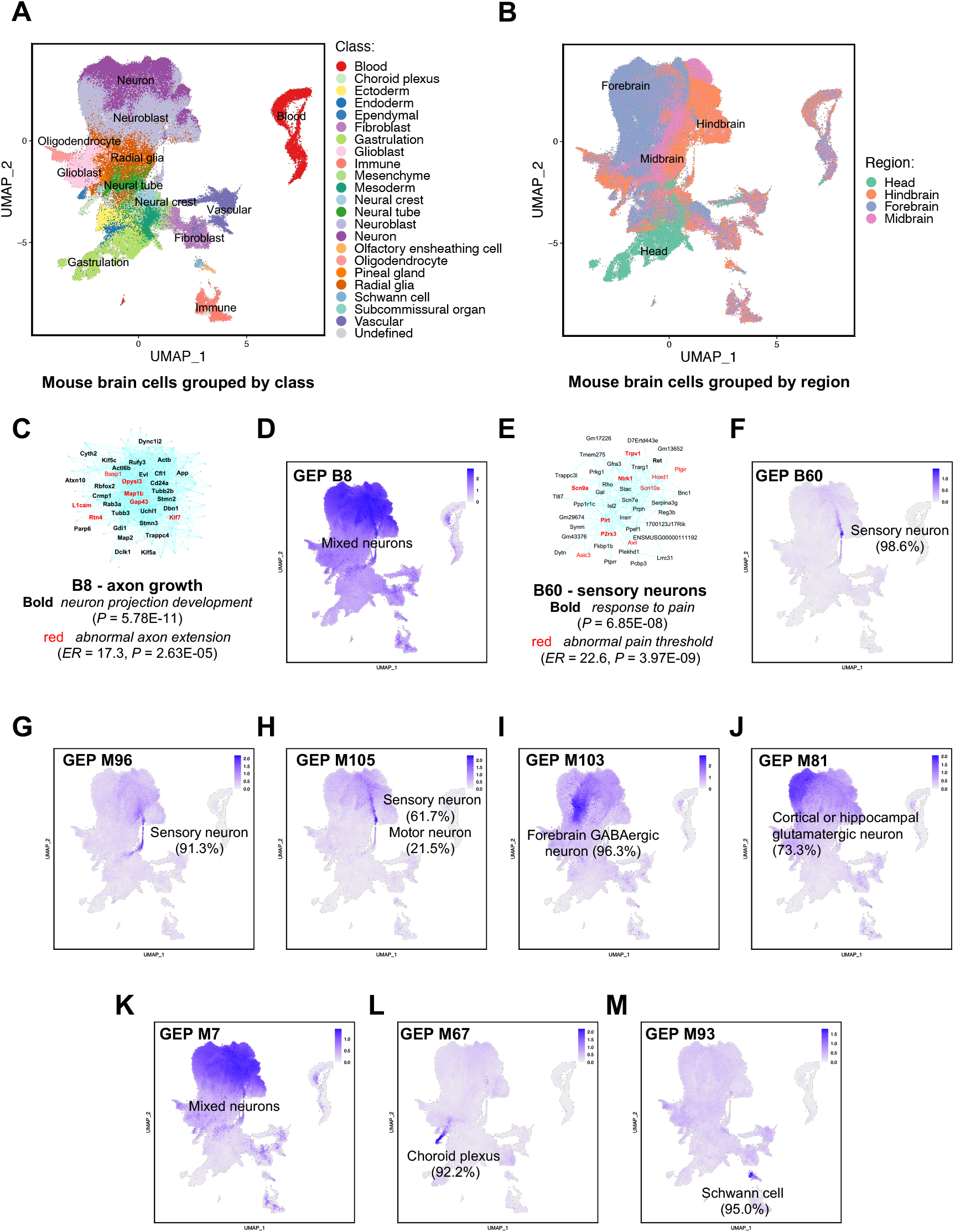
The GEPs identified from a mouse brain single-cell dataset. The dataset was obtained from (La Manno et al., 2021). Gene co-expression network analysis using SingleCellGGM identified 158 GEPs from the dataset. **(A)** A UMAP plot of mouse brain cells after dimension reduction via averageing-by-GEPs. Cells are colored according to cell class labels determined in the original study. **(B)** A UMAP plot of mouse brain cells colored according to their region of origins. **(C-D)** A subnetwork for GEP B8 and its expression across cells. The GEP is expressed across multiple types of neurons. **(E-F)** A subnetwork of GEP B60 and its expression across the cells. The number in (F) shows the percentage of cells belonging to the indicated cell type among all cells expressing the GEP, same as below. A GEP is considered to be expressed in a cell if its expression value is ≥ 1/2 of the GEP’s maximum expression value among all cells. GEP B60 is specifically expressed in sensory neurons. **(G-K)** Expression of MCA GEPs M7, M81, M96, M103, and M105 in the mouse brain dataset. Average expression values of the genes (ignoring those not contained in the brain dataset) within the GEPs are used to draw the expression map. **(L-M)** Expression of MCA GEPs M67 and M93 in the mouse brain dataset.

Our analysis identified GEPs which were broadly expressed in different types of neurons across diverse brain regions as well as GEPs specific to particular neurons. For example, GEP B8 is widely expressed in glutamatergic, GABAergic, and glycinergic neurons from the forebrain, midbrain, and hindbrain, and it is enriched with 32 genes significantly involved in *neuron projection development* (*P* = 5.78E-11; **Figure 5C, D**), including *Tubb3, Gap43*, and *Crmp1*, indicating that it is a common GEP used by different neurons for regulation of axon growth. Mutations of genes within the GEP result in *abnormal axon extension*, as well as *abnormal cerebral hemisphere, telencephalon*, and *temporal lobe morphologies* (*ER* = 17.3, 5.9, 5.0, and 6.8, *P* = 2.63E-05, 2.22E-07, 2.22E-07, and 1.40E-05, **Table S13**). Conversely, GEP B60 is specifically expressed within sensory neurons and enriched with 6 genes specifically involved in the *response to pain* (*P* = 6.85E-08), including *Nrtk1, Scn9a*, and *Trpv1* (**Figure 5E, F**). This GEP also contains 10 genes possessing the MP term *abnormal pain threshold*, representing a 22.6-fold enrichment compared to the background (*P* = 3.97E-09, **Table S13**). Thus, B60 is an important GEP for pain response.

Additional GEPs with specific expression in another group of sensory neurons (B52), cortical or hippocampal glutamatergic neurons (B1), forebrain GABAergic neurons (B21), midbrain GABAergic (B75) and dopaminergic (B85) neurons, hindbrain glycinergic neurons (B62) and serotoninergic neurons (B81), cerebellum glutamatergic neurons (B98), epithalamus glutamatergic neurons (B70), hypothalamus (B122), and Cajal-Retzius (B51) were also identified. These reveal transcriptional programs responsible for differentiating different types of neurons (**Figure S9**). Identity GEPs were also identified for other brain cell types or structures, such as glioblasts (B5), committed oligodendrocyte precursors (B25), Schwann cells (B39), optic cup (B121), lens (B149), Bergmann glia (B99), subcommissural organ hypendymal cells (B91), pia and meninges (B4), dura (B24), arachnoid (B40), extraembryonic endoderm (B6), cardiac mesoderm (B29), macrophage (B16), and microglia (B55) (**Figure S10**).

### GEPs facilitate universal cell type label transfer between single-cell studies

The GEPs identified by SingleCellGGM network analysis can be used to facilitate cell-type label transfer across different studies. Our analysis of MCA identified 5 GEPs (M7, M81, M96, M103, and M105) which represent different subtypes of neurons, but their subtype identities remain largely unknown. We mapped these GEPs onto the mouse brain dataset and uncovered that they have different expression patterns (**Figure 5G-K**). M96 is mainly expressed in sensory neurons, M105 in sensory and motor neurons, M103 in forebrain GABAergic neurons, and M81 in cortical or hippocampal glutamatergic neurons. M7, much like GEP B8, has expression throughout multiple types of neurons, and this GEP is particularly enriched with the GO term *neuro projection development* (*P* = 2.42E-17), indicating that it also regulates axon growth. Based on these results, we transferred these GEPs’ cell type labeling from the brain dataset to MCA, with M81, M96, M103, and M105 being marker GEPs for specific neurons and M7 being a general marker GEP for neurons.

Cell type annotations for GEPs can also be cross-referenced by multiple datasets. We found that annotations for two GEPs were different between the MCA and brain datasets. In MCA, cells annotated by M67 and M93 were labeled as Schwann cells and myocytes, respectively (**Figure 2A**). However, in the brain dataset, M67 is expressed in choroid plexus cells, while M93 is expressed in Schwann cells (**Figure 5L, M**). As M67 contains the choroid plexus marker genes *Ttr, Otx2, Kcne2*, and *Kcnj3* and M93 contains Schwann cell marker genes *Mpz, Sox10*, and *Gfra3*, we revised M67 and M93 to be marker GEPs for the choroid plexus and Schwann cells, respectively. Based on the marker genes they contain, we also refined or revised cell type annotations for another 13 MCA GEPs. For example, M17 was revised to be a marker GEP for ciliated cells and ependymal cells, M71 for alveolar epithelial type I (AT1) cells, and M112 for eosinophils (**Table S5**).

Finally, cell type labels of the GEPs can be transferred to novel single-cell datasets. We used the identity GEPs from MCA to annotate the TMS FACS dataset (Tabula Muris 2020). SingleCellGGM and UMAP analyses were conducted to generate a map for cells in TMS, which was then used to determine the expression of the MCA GEPs. In TMS, M1, a marker GEP for oligodendrocytes, is expressed across a group of cells that are clustered together, and they were thus annotated as oligodendrocytes (**Figure 6A**). Supporting this annotation, the oligodendrocyte marker genes *Mog* and *Olig1* had similar expression patterns as GEP M1 (**Figure 6B-C**). In contrast, the expression map of M26, a GEP for 3T3 cells, contained scattered and low-expression loci (**Figure 6D**). We considered that M26 was not expressed and that no 3T3 cells were found in TMS, consistent with the fact that this cell line was not included in the original TMS sample. Based on these observations, we developed an algorithm named <u>Cel</u>l labeling via <u>g</u>ene <u>e</u>xpression <u>p</u>rograms (CellGEP) to facilitate cell label transfer between datasets. CellGEP first determines which GEPs are expressed in a dataset and uses the expressed GEPs to annotate its cells accordingly (see Materials & Methods for details). In total, 94,582 cells in TMS were annotated by 70 MCA GEPs, while 16,242 cells remained unannotated (**Figure 6E**). Similar to M1, other GEPs contain known marker genes with expression patterns similar to the GEPs themselves, which supports the assignment of labels based on GEPs (**Figure S11A**). We also used Harmony (Korsunsky et al. 2019) to integrate the MCA and TMS datasets and uncovered that after integration, cells annotated by the same GEPs in MCA and TMS all tended to locate in overlapping spaces, indicating that the GEPs annotate the same cell types in both datasets (**Figure S11B**). Of the 70 GEPs, 68 were supported by marker gene expression and/or by co-localization across the integrated cell space (**Figure S11C**). The remaining two GEPs, M56 and M85, annotated 1 and 7 cells, respectively. Additionally, CellGEP supports the identification of novel cell types. The unannotated cells form their own cell clusters in the TMS cell map. They could represent novel cell types not present in MCA, and an analysis on the TMS dataset should reveal their corresponding GEPs.

**Figure 6.**
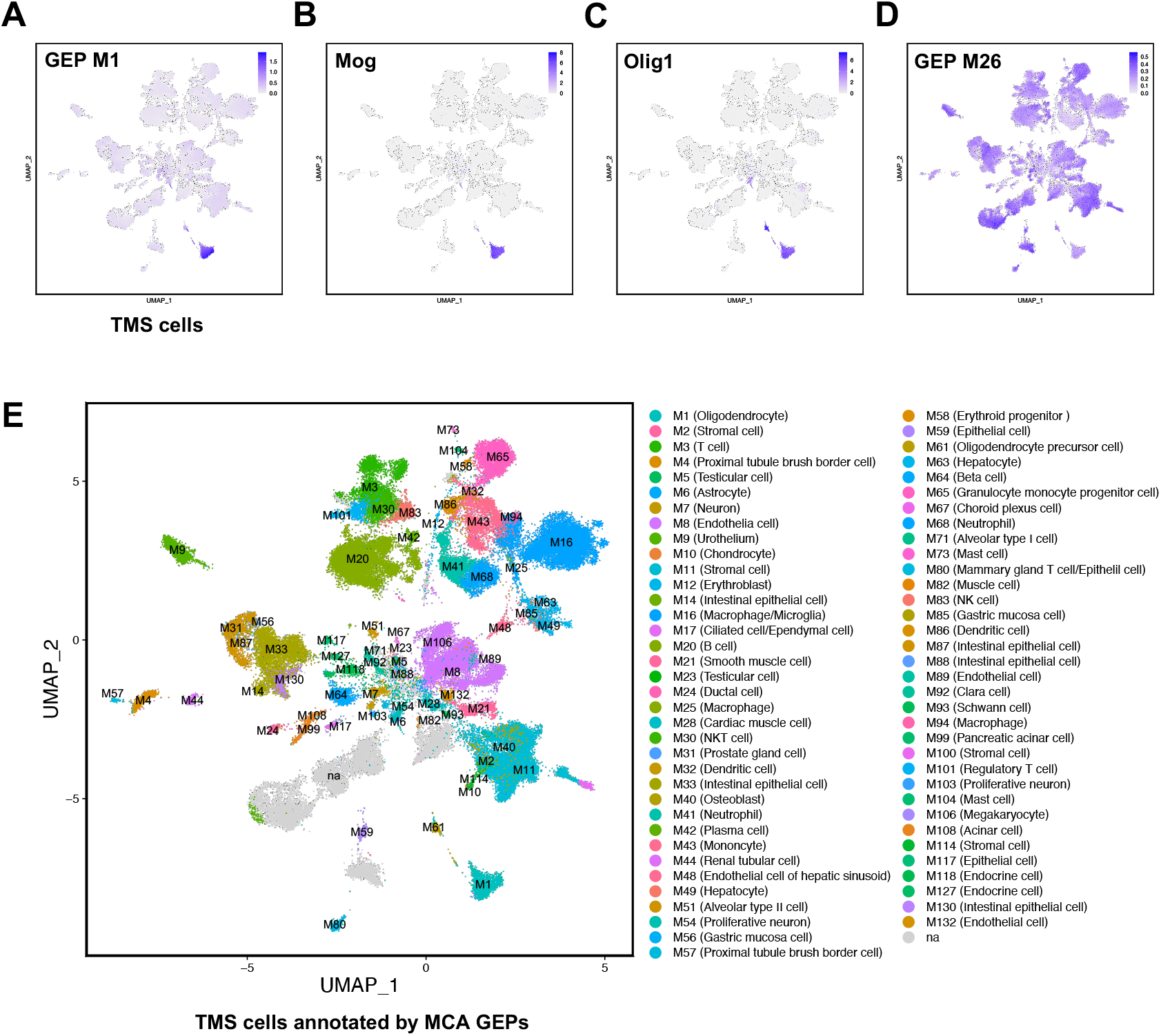
Cell type label transfer between the MCA and TMS datasets. **(A-C)** Expression of MCA GEP M1, a marker GEP for oligodendrocytes, and *Mog* and *Olig1*, two oligodendrocyte marker genes, in TMS cells. **(D)** Expression of MCA GEP M26, a marker GEP for 3T3 cells, in TMS cells. **(E)** TMS cells annotated by MCA GEPs. MCA GEPs are used to annotate the TMS cells via CellGEP algorithm. Cells are colored according to their GEP labels.

## Discussion

We developed a SingleCellGGM algorithm to conduct GCN analysis on single-cell transcriptome datasets. Single-cell GCN analysis is challenging due to excessive dropout expression values. Previous methods have used cell grouping or gene imputation to address the problem of dropouts (Li and Li 2018; van Dijk et al. 2018; Baran et al. 2019; Xu et al. 2022a), which still may cause loss of cell resolution or introduce spurious correlations. SingleCellGGM does not require cell grouping or gene imputation but still achieves satisfactory GCN results. When applied to the MCA and mouse brain datasets, SingleCellGGM generated two GGM gene co-expression networks from which GEPs of biological significance were identified. Many of the GEPs have significantly enriched GO and MP terms, and a portion of them have cell-type specific expression. The GEPs also enabled for development of a dimension reduction method via averaging-by-GEPs for single-cell transcriptome analysis, which generated UMAP plots that clearly separated the different cell types in both the MCA and mouse brain datasets (**Figure 1H and Figure 5A**). In addition to identifying GEPs of biological significance, SingleCellGGM also has the capacity to handle large-scale transcriptomes, including the mouse brain dataset with 292,495 cells. Therefore, we envision that SingleCellGGM may also be easily applied to analysis of other single-cell transcriptome datasets and achieve similar performance.

The GEPs identified by SingleCellGGM network analysis provide an ample number of candidate genes to study cellular functions. Based on their expression and functional annotations, we divided the GEPs into identity, activity, and morphogenesis GEPs. The concepts of identity and activity GEPs were proposed in a previous study (Kotliar et al. 2019), while morphogenesis GEPs are derived from our current analysis. Most identity GEPs have cell-type specific expression and are enriched with genes required for the functions of the corresponding cells, such as the GEPs for osteoblasts (M40), NK cells (M83), and sensory neurons (B60) (**Table S6, S13**). These GEPs cover a broad range of cell types and provide numerous candidate genes to study the corresponding cellular functions. Correspondingly, activity and morphogenesis GEPs usually have shared expression across different cell types, including GEPs involved in cell-cycle regulation (M18), general stress response (M53), and tissue development (M72). These GEPs are also enriched with genes functioning in aligned biological processes. Notably, these GEPs have been largely overlooked by previous single-cell studies, potentially due to previous studies mainly focusing on identifying marker genes specific for individual cell types. However, these GEPs also contain important cellular regulators. For example, morphogenesis GEPs in both MCA and the brain dataset may play important roles in cellular development and tissue patterning. Our SingleCellGGM analysis revealed transcriptional programs that have been overlooked by previous analyses.

GEPs may also be used to annotate individual mouse cells directly. As demonstrated above, we can assign marker GEPs to cells in MCA based on their expression. Most marker GEPs have cell type specificities, and can be used to annotate cells for their cell types (**Figure 2A**). The GEPs usually contain multiple marker genes aligned to corresponding cell types, and marker gene expression often matches that of the GEPs themselves, providing further support for GEP-based cell type annotation. Additionally, GEPs are also conserved across datasets which facilitates label transfer between different single-cell studies. We uncovered that five neuron-related GEPs from MCA had specific expression across different subtypes of neurons within the mouse brain dataset, and we thus transferred these neuron subtype labels from the brain dataset to the MCA dataset. The GEPs derived from the MCA dataset were also used to annotate and label cells in the TMS FACS dataset (**Figure 6**). Such label transferring was supported by two observations: first, within the TMS dataset, MCA GEPs shared similar expression patterns to the known marker genes they contain; and second, the cells annotated by the same GEPs from both MCA and TMS co-localized in similar cell spaces after the two datasets were integrated via Harmony. Therefore, the GEPs can be considered as transcriptional signatures of different cell types, and can be used as marker GEPs across single-cell studies to annotate cells. In the future, similar analyses should be conducted on other single-cell transcriptome datasets to identify novel marker GEPs for cell types not covered in the current study.

In conclusion, we have developed a SingleCellGGM algorithm to conduct gene co-expression network analysis for single-cell transcriptomes. The GEPs identified from the resulted networks can be used to facilitate gene function studies across different cell types. They can also be used to annotate individual cells directly, facilitating universal cell type label transfer between different single-cell studies. Our approach thus provides a unique GEP-based perspective for analysis of single-cell transcriptome datasets.

## Materials and Methods

### Single-cell gene co-expression network analysis using SingleCellGGM

A SingleCellGGM algorithm was developed to conduct single-cell gene co-expression network analysis. The algorithm was modified from a GGM algorithm developed previously for gene co-expression analysis in bulk transcriptome datasets (Ma et al. 2007). The original algorithm was developed mainly using the R language for *pcors* calculation, and a modified MATLAB version of the algorithm, called rgsGGM and also for analyzing bulk datasets, was developed recently (Wang et al. 2023). We developed SingleCellGGM based on the MATLAB version of the algorithm. Briefly, SingleCellGGM uses a process consisting of ∼20,000 iterations to calculate *pcors* between genes. In each iteration, 2,000 genes are randomly selected, and a covariance matrix Σ for the selected genes is estimated via the sample covariance estimator based on gene expression values. Note that the original GGM algorithm uses a shrinkage approach for estimating the covariance matrix (Schafer and Strimmer 2005), in order to accommodate the “small n, large p” scenario where the number of bulk transcriptome samples (n) was smaller or not much larger than the number of selected genes (p). For single-cell samples, the number of cells is usually much larger than the number of selected genes, therefore we choose to use the sample covariance estimator to estimate the covariance matrix directly. The estimated covariance matrix is then used to calculate *pcors* between selected genes, which is related to the inverse of the covariance matrix Σ (Whittaker 1990):

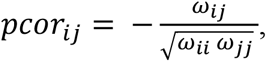

where,

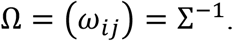

After 20,000 iterations, every gene pair is sampled over multiple iterations and their *pcor* is calculated multiple times. The *pcor* with the lowest absolute value is then chosen as the final *pcor* of that gene pair. For every gene pair, SingleCellGGM also calculates how many cells co-expression occurs in, meaning both genes have nonzero expression values in those cells. The gene pairs with *pcors* ≥ a selected cutoff value (e.g., 0.03) and co-expressed in ≥ a selected number of cells (e.g., 10) are chosen to construct a final GGM gene co-expression network. A MATLAB script (SingleCellGGM.m) was developed to conduct the analysis.

To conduct gene co-expression network analysis for a single-cell transcriptome dataset, its gene count matrix was first normalized and log-transformed via log_2_(CP10K + 1). The genes expressed in fewer than a selected number of cells were filtered out, and the log-transformed matrix of the remaining genes was then used for *pcors* calculation via SingleCellGGM. The resulting network was clustered into gene co-expression modules via MCL clustering algorithm (Enright et al. 2002), and the modules were considered as GEPs.

### Dimension reduction via averaging-by-GEPs

For a single-cell transcriptome dataset, we first used its gene count matrix to create a single-cell analysis object in Seurat (v4) (Hao et al. 2021). We then conducted a gene co-expression network analysis using SingleCellGGM to identify GEPs. Based on the identified GEPs, we used the normalized and log-transformed expression matrix of the dataset to calculate average expression values of the genes within the GEPs in every cell to obtain a GEP expression matrix. The GEP expression matrix was scaled by GEPs and used to create a dimension reduction object via the CerateDimReducObject function in the SeuratObject package in R, which was further used for UMAP analysis and cell map visualization.

### Single-cell and bulk transcriptome datasets used in the analysis

The count matrix of the MCA dataset (“MCA_Figure2-batch-removed.txt.tar.gz”) was downloaded as provided by the original study (https://figshare.com/s/865e694ad06d5857db4b) (Han et al. 2018), together with cell type annotations for the cells. The dataset, described in Figure 2 of the original publication, was subsampled from the complete MCA dataset. We selected 24,802 genes with Ensembl gene IDs to conduct gene co-expression network analysis via SingleCellGGM. The gene pairs with *pcors* ≥0.03 and co-expressed in ≥10 cells were chosen to construct a gene co-expression network named MCA_GGM. The network was clustered by the MCL algorithm with the parameters “-I 1.7 -scheme 7”, and 137 gene co-expression modules with ≥10 genes were obtained, which were considered GEPs. Using these GEPs, a dimension reduction for the MCA dataset by averaging-by-GEP was conducted, followed by a UMAP analysis with the parameter “spread = 0.3” in Seurat.

The mouse brain dataset was downloaded from the Mouse Brain Atlas database (http://mousebrain.org/development/downloads.html) (La Manno et al. 2021). The downloaded file (“dev_all.loom”) contained expression values for 31,053 genes in 292,495 cells, as well as annotations for all cells. We chose 24,080 genes expressed in ≥10 cells for SingleCellGGM and downstream analysis, following a procedure akin to the analysis for the MCA dataset.

We also downloaded the TMS FACS single-cell dataset from the CZ CELLxGENE Discover database (https://cellxgene.cziscience.com/collections/0b9d8a04-bb9d-44da-aa27-705bb65b54eb). The dataset contained gene count expression values for 21,069 genes in 110,824 cells. We chose 20,911 genes expressed in ≥10 cells to construct a gene co-expression network named TMS_GGM via SingleCellGGM. We obtained mouse bulk RNA-seq transcriptome datasets from the ARCHS4 (v10) database (Lachmann et al. 2018). After quality control and manual removal of single-cell transcriptome samples, transcriptome data from 91,658 mouse bulk RNA-seq were used to construct a GGM gene co-expression network named ARCHS4_GGM. We used a process much like SingleCellGGM to construct a GGM network for bulk transcriptomes, with the difference being that gene expression data were log-normalized via log_2_(CPM +1) and gene pairs were not required to be co-expressed in more than a certain number of samples. For every GEP identified from MCA_GGM, we used its genes to extract two subnetworks from TMS_GGM and ARCHS4_GGM. The RBGL package in R was used to identify the maximum connected component contained within the subnetworks. The number of genes contained within the maximum connected components were compared with the number of genes contained in the original GEP to ultimately evaluate the robustness of the GEP.

### Functional annotation of the identified GEPs

We obtained GO annotations for mouse genes from Ensembl BioMart on June 20, 2022, and MP annotations (“MGI_PhenotypicAllele.rpt”) from the MGI database on October 18, 2022. The annotations were used for GO and MP enrichment analysis for GEPs via a hypergeometric test. We obtained known marker genes for mouse cell types from the CellMarker and CellTaxonomy databases as well as the original MCA study (Han et al. 2018; Jiang et al. 2022; Hu et al. 2023).

### Marker GEP identification and cell type label transfer

We calculated average gene expression values for every identity GEP within individual MCA cells. A GEP was considered to be expressed in a cell if the GEP’s expression value ≥ 1/2 of the maximum GEP expression value among all cells. Across all expressed identity GEPs in a cell, the one with the highest expression value was considered to be the marker GEP. Only a portion of cells are annotated in this initial step. The average GEP expression values of the cells were also used to construct a K-nearest neighbor (KNN) network to identify cell neighbors. If none of the neighbors of a cell shared the same marker GEP with that cell, we reassigned the cell as unannotated. We then used a neighboring voting procedure to propagate marker GEP labels to unannotated cells. Briefly, if an unannotated cell was connected to three or more cells with the same label and the unannotated cell’s corresponding GEP’s expression level was ≥ 0.3, the label was then assigned to the unannotated cell. This step was repeated three times to propagate labels to neighboring cells.

We developed a CellGEP algorithm (‘cellgep.R’) in R to facilitate cell type label transfer between single-cell datasets. For a set of selected identity GEPs, CellGEP first removed the genes not contained in an unannotated target dataset, and only kept the GEPs with 10 or more genes for analysis. CellGEP next determined which GEPs are expressed within the target dataset. A GEP is considered to be expressed in the dataset if, at a cutoff level between 50% and 100% of its maximal expression level among all cells within the dataset, at least 98% of the cells expressing the GEP are connected as a single connected group or 100% of them form at most two connected groups. CellGEP used a dynamic process to determine which cutoff value satisfied these criteria and covered the most cells. CellGEP then used the cutoff value to determine which cells expressed the GEP in the target dataset. The GEPs that did not meet the criteria were considered not expressed in the target dataset. Once the expressing identity GEPs were determined for all cells, a neighboring voting step was repeated three times to propagate cell labels to neighboring unannotated cells.

### Comparing SingleCellGGM to other tools

We used the ppcor and SILGGM algorithms to analyze the same log-transformed MCA gene expression matrix analyzed by SingleCellGGM. We then selected the top 127,229 co-expressed gene pairs according to *pcors* calculated in both algorithms and constructed two co-expression networks named MCA_ppcor and MCA_SILGGM. These two networks were clustered via MCL into gene co-expression modules with the same parameters as MCA_GGM, and the modules were considered GEPs. The GEPs identified by SingleCellGGM, ppcor, and SILGGM were compared to determine which method identified the most genes in GEPs with significant enrichment GO terms.

We used the cNMF method to analyze the MCA dataset, with the factor number set as 137, the same as the number of GEPs identified by SingleCellGGM. We then determined the cell type specificities of the programs identified by cNMF and compared them to GEPs identified by SingleCellGGM to determine how many identity or activity GEPs were identified by each method.

## Data and Code Availability

The mouse single-cell datasets were obtained from public databases as described in the text. The SingleCellGGM and CellGEP algorithms are available from GitHub (https://github.com/MaShisongLab/SingleCellGGM and https://github.com/MaShisongLab/CellGEP).

## CRediT Authorship Contribution Statement

**Yupu Xu:** Methodology, Investigation, Formal analysis, Writing – Original draft. **Yuzhou Wang:** Formal analysis. **Shisong Ma:** Conceptualization, Methodology, Formal analysis, Supervision, Writing – Original draft, Review & Editing, Funding acquisition.

## Acknowledgements

We thank the USTC Supercomputing Center and USTC School of Life Sciences Bioinformatics Center for providing the computing resources. This work was supported by grants from the Strategic Priority Research Program of the Chinese Academy of Sciences (XDA24010303), the National Natural Science Foundation of China (31770268), the Fundamental Research Funds for the Central Universities (WK2070000091), and University of Science and Technology of China (Start-up fund to S.M.).

## Supplemental Information

**Figure S1. Distribution of all *pcors* calculated using the MCA dataset**. Most of the gene pairs have *pcors* values ranging between -0.01 and 0.01, indicating no correlation between those gene pairs. The gene pairs with *pcors* ≥ 0.03 are located at the far-right end of the distribution curve, indicating a very stringent cutoff value.

**Figure S2. Comparison between MCA_GGM and TMS_GGM. (A)** A subnetwork extracted from TMS_GGM of the 179 genes within GEP M8 of MCA_GGM. The largest connected component contains 147 genes, or 82% of the 179 genes within GEP M8. **(B)** A histogram showing the module conservation between TMS_GGM and MCA_GGM.

**Figure S3. A heatmap showing the cell-type-specific expression of genes within the GEPs**. Thirty genes were randomly selected from the GEPs, and thirty cells were randomly sampled from MCA according to their cell type labeling in the original study (Han et al., 2018).

**Figure S4. Tissue origins of the cells annotated by GEP M104**.

**Figure S5. Expression of GEP M35 and M53 in MCA cells. (A)** Expression of GEP M35 in MCA cells. **(B)** Expression of GEP M53 in MCA cells.

**Figure S6. Expression of morphogenesis GEPs in MCA cells**. Expression of GEPs M70, M72, M77, M95, and M137 in MCA cells as grouped by tissues (A, C, E, G, I) or cell types (B, D, F, H, J).

**Figure S7. Cell-type-specific usage of the GEPs identified by cNMF from the MCA dataset. (A)** Expression maps showing the usage levels of GEP C1 – C10 identified by cNMF. Only 10 GEPs are shown due to limited space. These GEPs display specific usage in one or a limited number of related cell types. **(B)**An expression map showing the usage level of GEP C84. This is the only GEP that displays shared usages across several different cell types. The color scale for each plot represents the raw usage level of the indicated GEPs. The max cutoff value of usage levels are set at 200 when drawing these expression maps.

**Figure S8. A UMAP plot of cells within the mouse brain dataset**. The cells are colored according to their subclass label annotated in the original study (La Manno et al., 2021).

**Figure S9. Identity GEPs with specific expression in various types of neurons**.

**Figure S10. Identity GEPs with specific expression in various brain cell types or structures**.

**Figure S11. Two lines of evidence supporting cell type label transfer between the MCA and TMS datasets. (A)** Expression of MCA GEPs and known marker genes contained in TMS cells. The left column shows the expression of marker GEPs, and the right three columns to show the expression of known marker genes contained within the GEP. Most GEP expression patterns are similar to the known marker genes. Only GEP M2 – M10 are shown due to limited space. **(B)** The cells annotated by the same GEPs from MCA and TMS after the two datasets were integrated. The cells annotated by the same GEPs from these two datasets often overlap with each other after integration. Only GEP M1-M10 are shown due to limited space. **(C)** Evidence supporting GEP identifying the same cell types in both datasets.

**Table S1. Co-expressed gene pairs used for MCA_GGM gene co-expression network construction. Table S2. The 137 GEPs identified from MCA_GGM**.

**Table S3. GO analysis results of the GEPs in MCA_GGM**.

**Table S4. Conservation of the MCA GEPs in the ARCHS4 and TMS datasets. Table S5. Three categories of GEPs identified from MCA_GGM**.

**Table S6. MP analysis results of the GEPs in MCA_GGM**.

**Table S7. Co-expression networks constructed for the MCA dataset using SILGGM and ppcor. Table S8. A comparison of the GEPs identified from MCA by three different methods**.

**Table S9. Co-expressed gene pairs used for MouseBrain_GGM gene co-expression network construction**.

**Table S10. The 158 GEPs identified from MouseBrain_GGM. Table S11. GO analysis results of the GEPs in MouseBrain_GGM**.

**Table S12. Three categories of GEPs identified from MouseBrain_GGM. Table S13. MP analysis results of the GEPs in MouseBrain_GGM**.

