## Supplemental Figures for "SingleCellGGM enables gene expression program identification from single-cell transcriptomes and facilitates universal cell label transfer"

### Figure S1

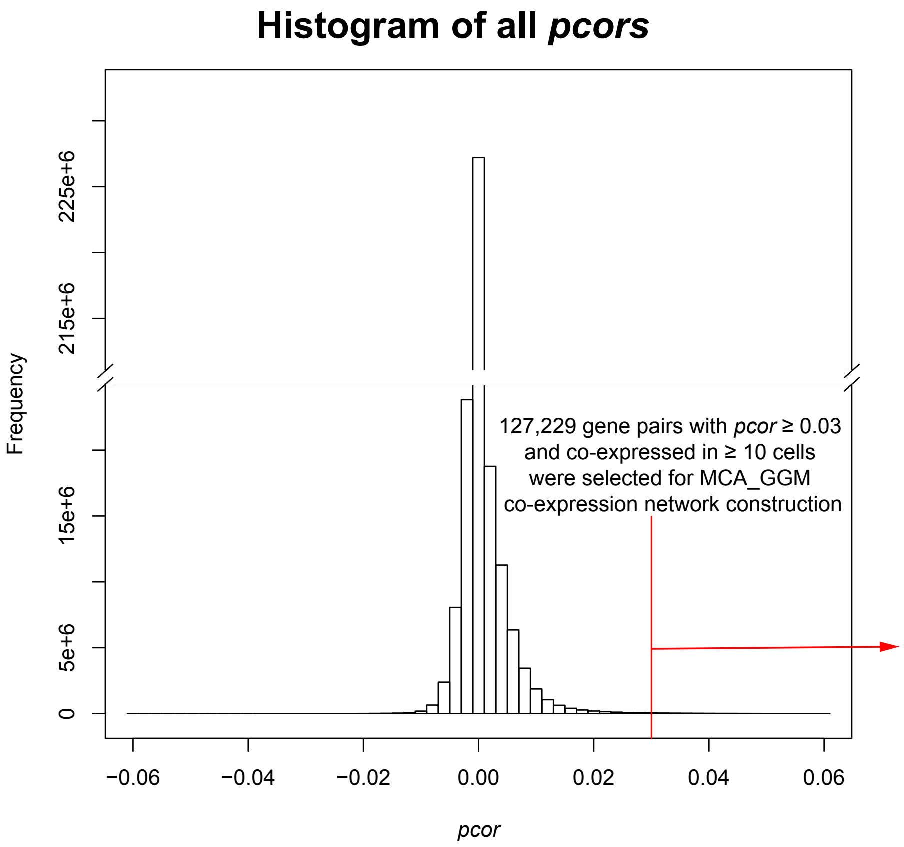


**Figure S1. Distribution of all *pcors* calculated using the MCA dataset.** Most of the gene pairs have *pcors* values ranging between -0.01 and 0.01, indicating no correlation between those gene pairs. The gene pairs with *pcors* ≥ 0.03 are located at the far-right end of the distribution curve, indicating a very stringent cutoff value.

**Figure S2**

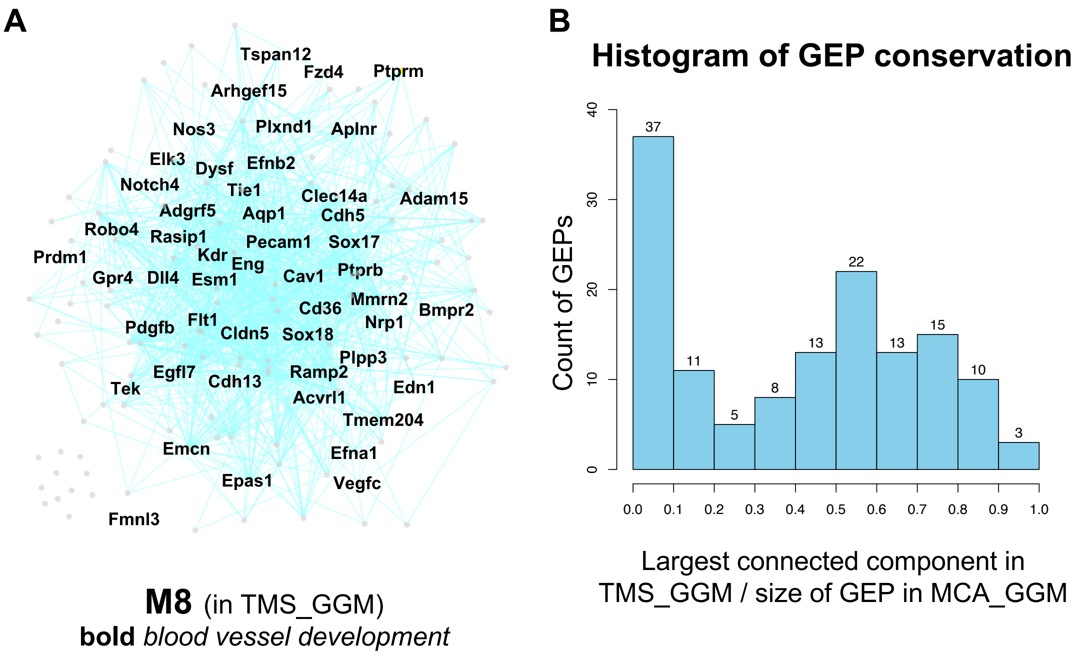


**Figure S2. Comparison between MCA_GGM and TMS_GGM. (A)** A subnetwork extracted from TMS_GGM of the 179 genes within GEP M8 of MCA_GGM. The largest connected component contains 147 genes, or 82% of the 179 genes within GEP M8. **(B)** A histogram showing the module conservation between TMS_GGM and MCA_GGM.

**Figure S3**

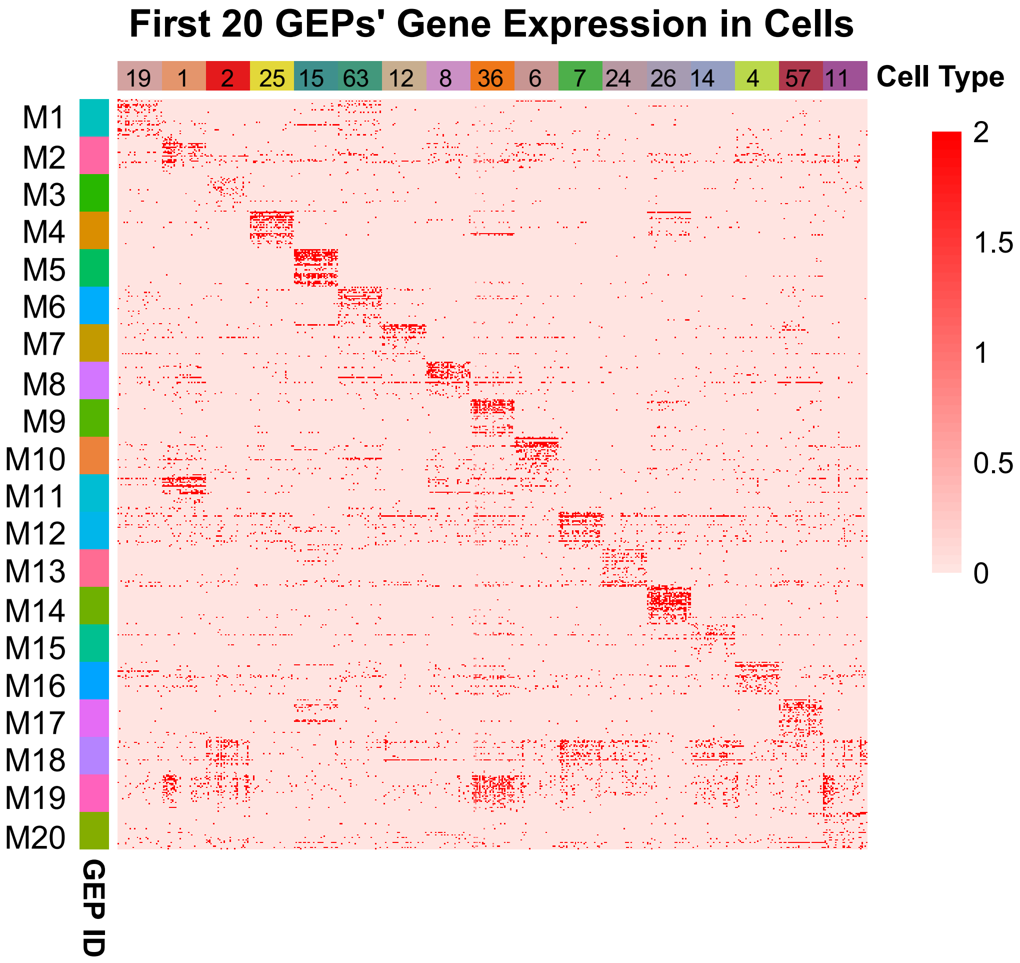


**Figure S3. A heatmap showing the cell-type-specific expression of genes within the GEPs.** Thirty genes were randomly selected from the GEPs, and thirty cells were randomly sampled from MCA according to their cell type labeling in the original study (Han et al., 2018).

### Figure S4

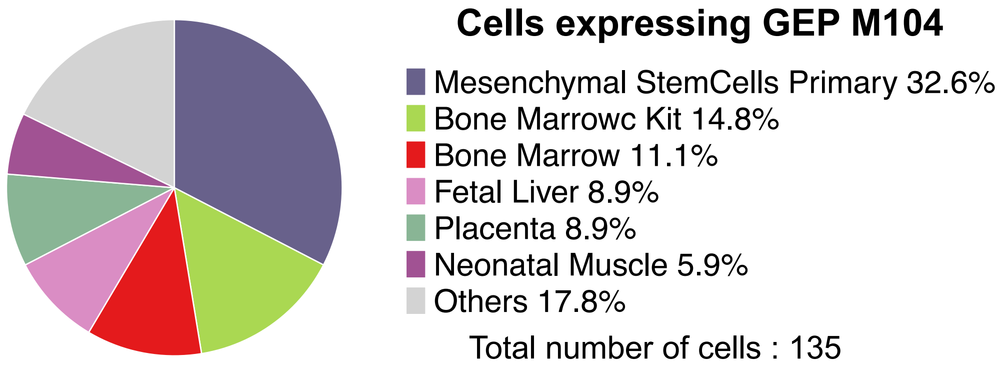


**Figure S4. Tissue origins of the cells annotated by GEP M104.**

### Figure S5

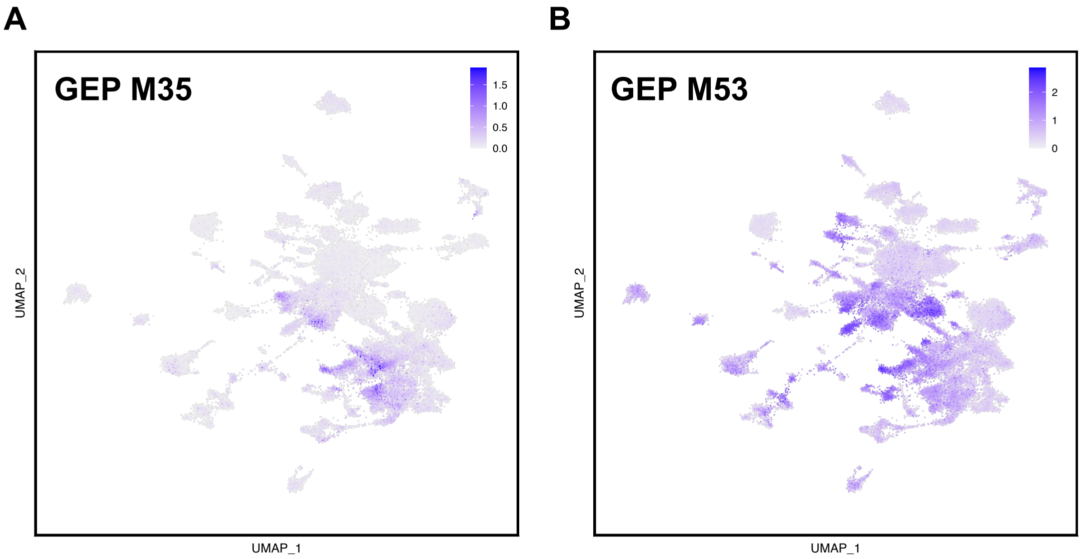


**Figure S5. Expression of GEP M35 and M53 in MCA cells. (A)** Expression of GEP M35 in MCA cells. **(B)** Expression of GEP M53 in MCA cells.

### Figure S6

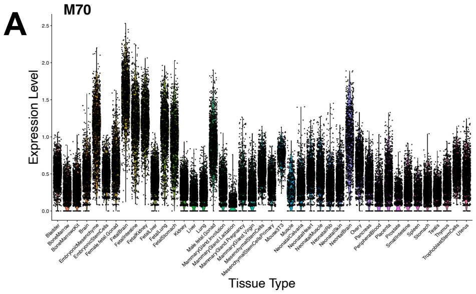


**Expression of GEP M70 grouped by tissues**

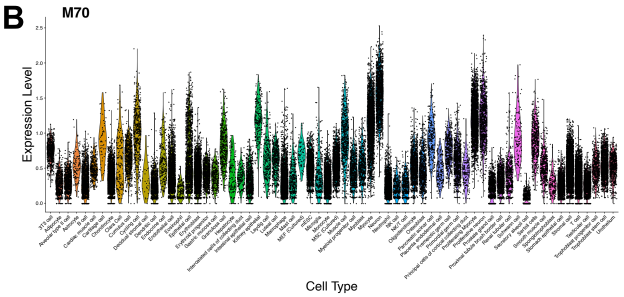


**Expression of GEP M70 grouped by cell types**

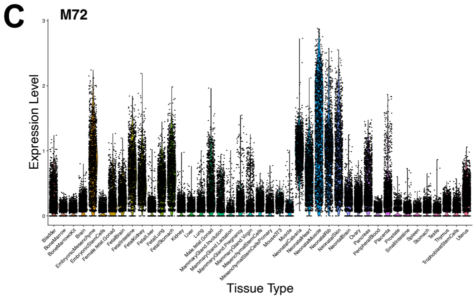


**Expression of GEP M72 grouped by tissues**

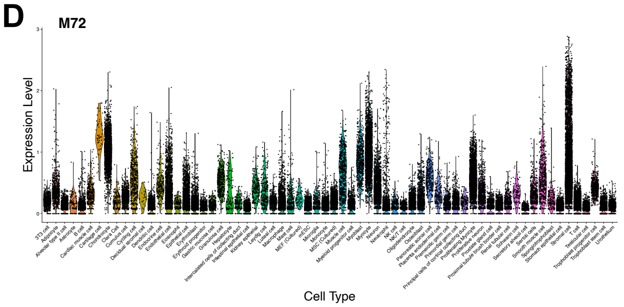


**Expression of GEP M72 grouped by cell types**

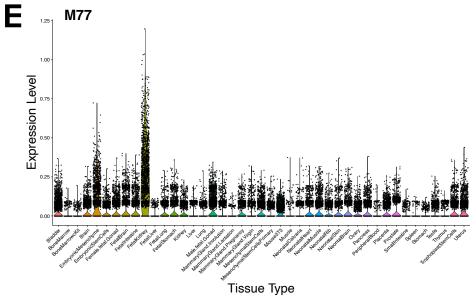


**Expression of GEP M77 grouped by tissues**

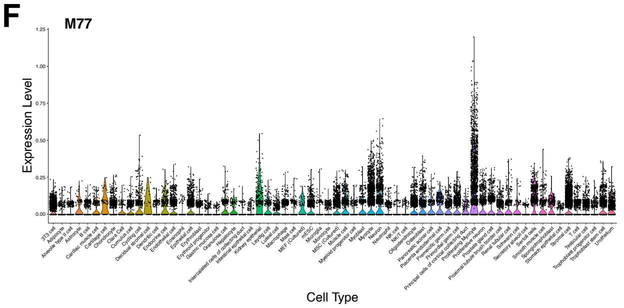


**Expression of GEP M77 grouped by cell types**

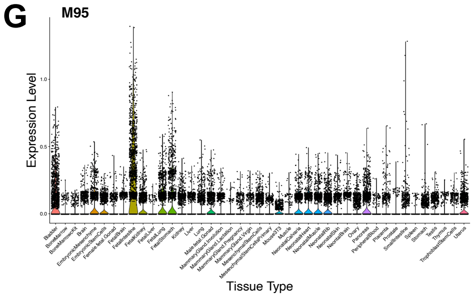


**Expression of GEP M95 grouped by tissues**

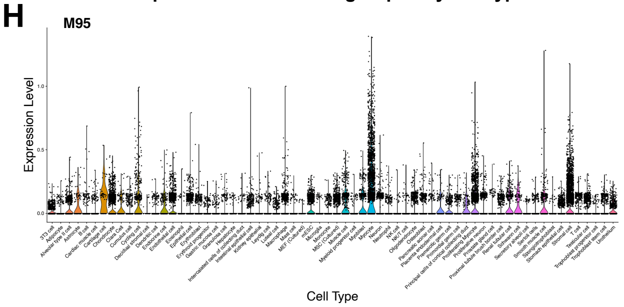


**Expression of GEP M95 grouped by cell types**

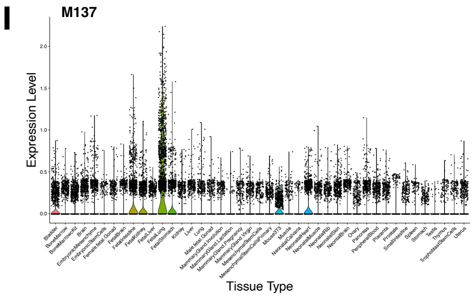


**Expression of GEP M137 grouped by tissues**

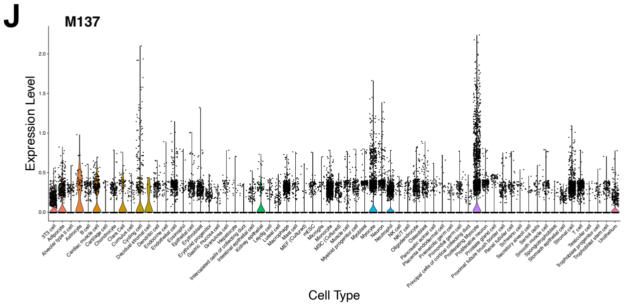


**Expression of GEP M137 grouped by cell types**

**Figure S6. Expression of morphogenesis GEPs in MCA cells.** Expression of GEPs M70, M72, M77, M95, and M137 in MCA cells as grouped by tissues (A, C, E, G, I) or cell types (B, D, F, H, J).

### Figure S7

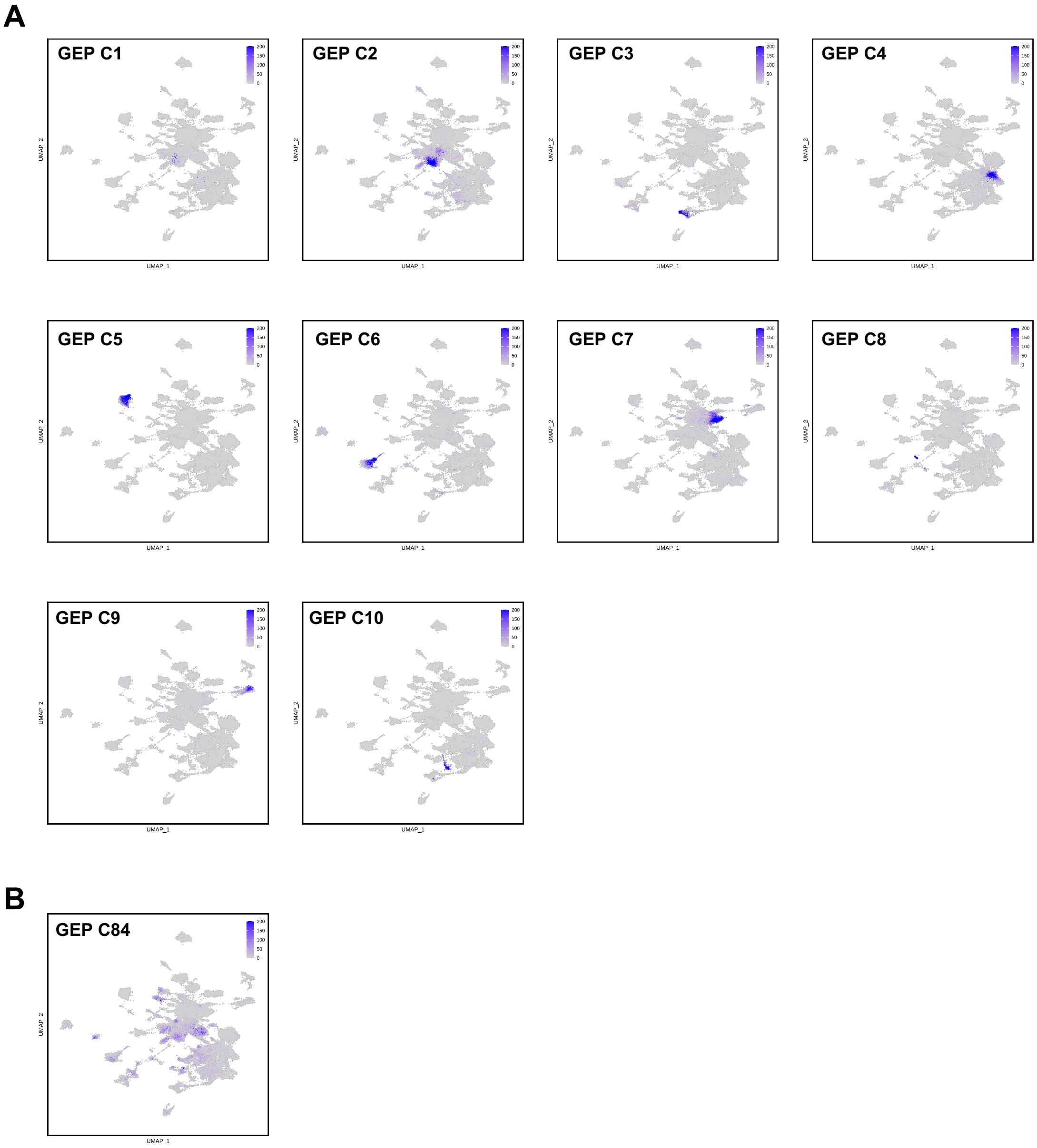


**Figure S7. Cell-type-specific usage of the GEPs identified by cNMF from the MCA dataset. (A)** Expression maps showing the usage levels of GEP C1 – C10 identified by cNMF. Only 10 GEPs are shown due to limited space. These GEPs display specific usage in one or a limited number of related cell types. **(B)** An expression map showing the usage level of GEP C84. This is the only GEP that displays shared usages across several different cell types. The color scale for each plot represents the raw usage level of the indicated GEPs. The max cutoff value of usage levels are set at 200 when drawing these expression maps.

### Figure S8

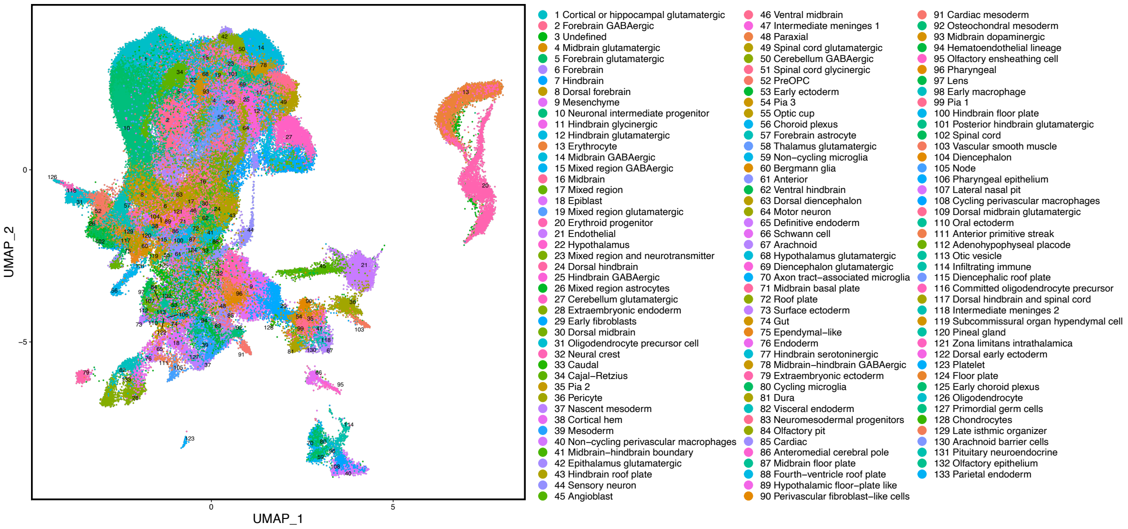


**Mouse brain cells grouped by subclass**

**Figure S8. A UMAP plot of cells within the mouse brain dataset.** The cells are colored according to their subclass label annotated in the original study (La Manno et al., 2021).

### Figure S9

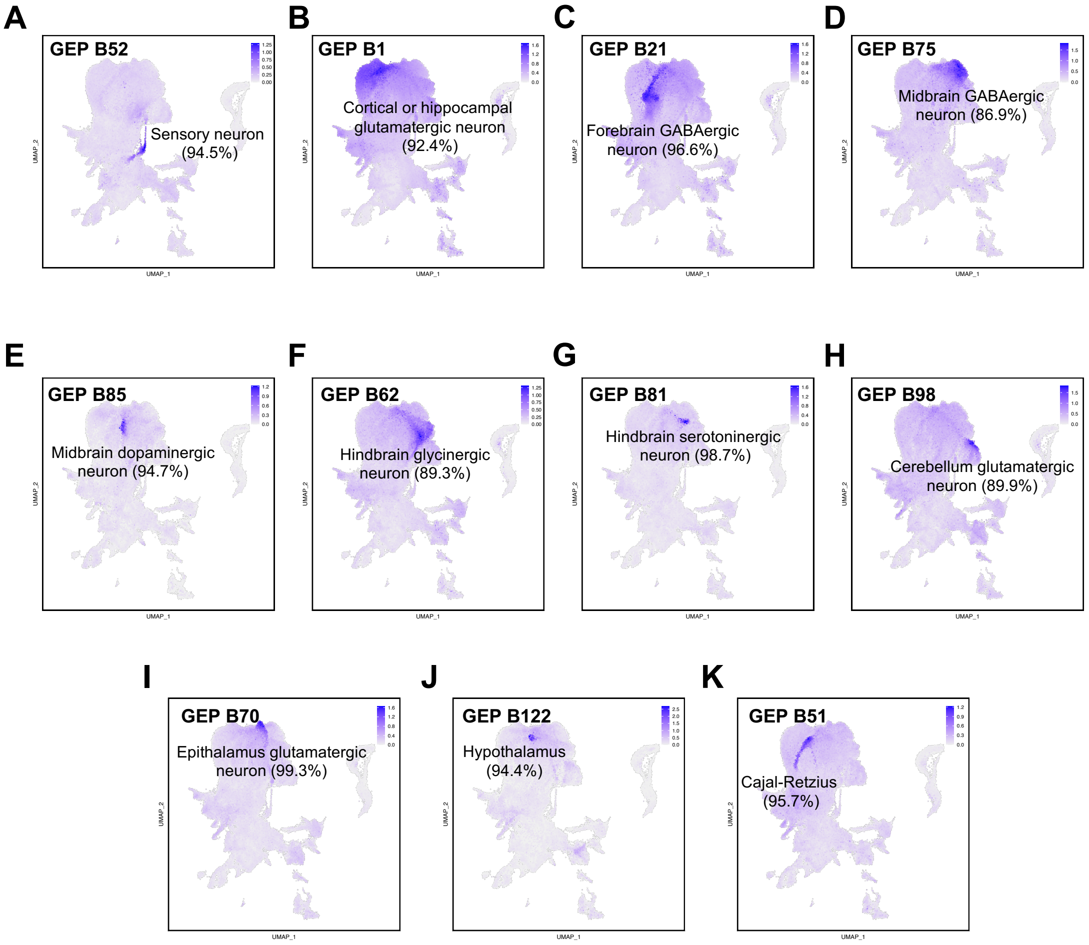


**Figure S9. Identity GEPs with specific expression in various types of neurons.**

### Figure S10

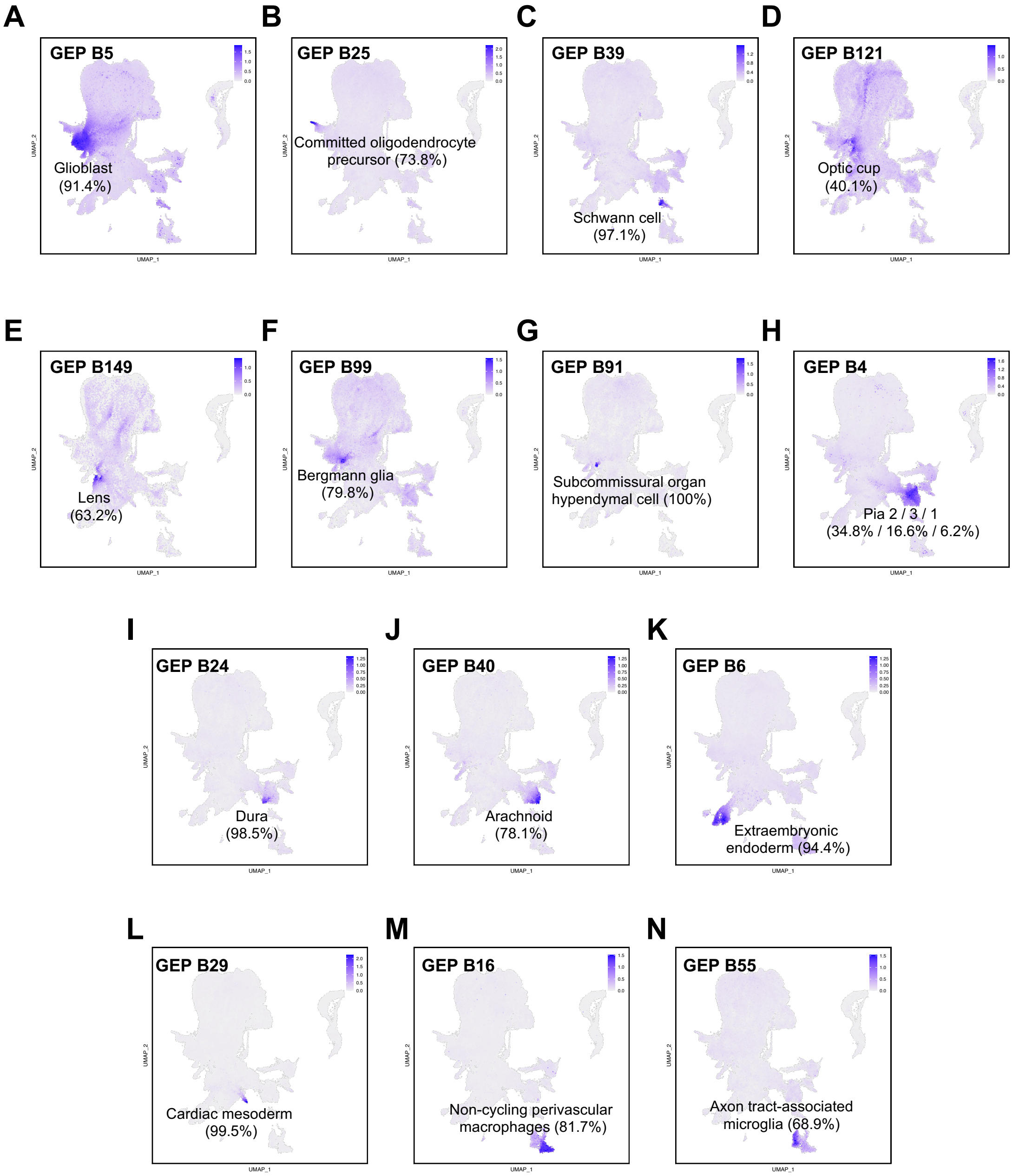


**Figure S10. Identity GEPs with specific expression in various brain cell types or structures.**

### Figure S11

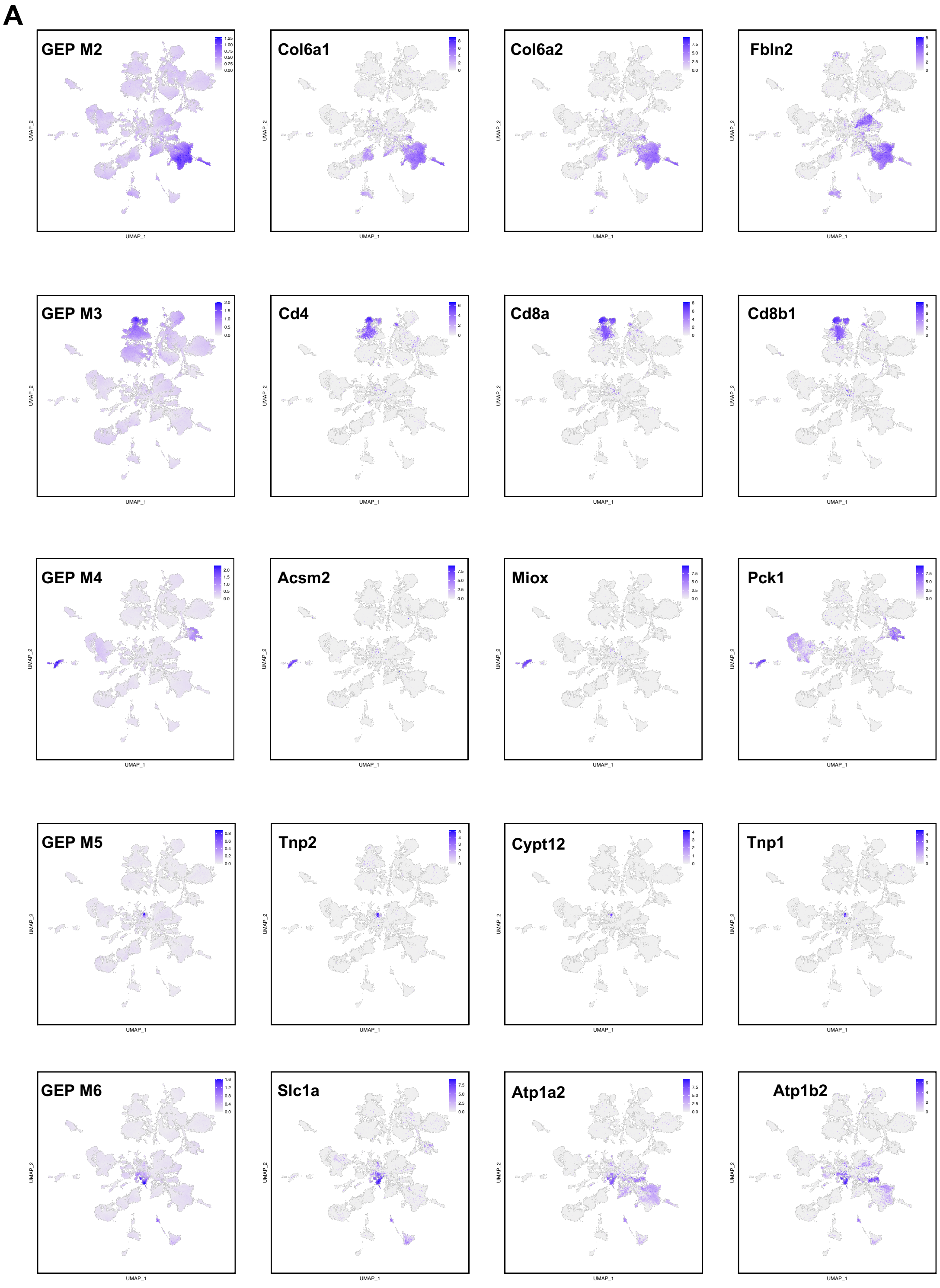
A
GEP M2
Col6a1
Col6a2
Fbln2
GEP M3
Cd4
Cd8a
Cd8b1
GEP M4
Acsm2
Miox
Pck1
GEP M5
Tnp2
Cypt12
Tnp1
GEP M6
Slc1a
Atp1a2
Atp1b2
UMAP_1
UMAP_2

### Figure S11 continued

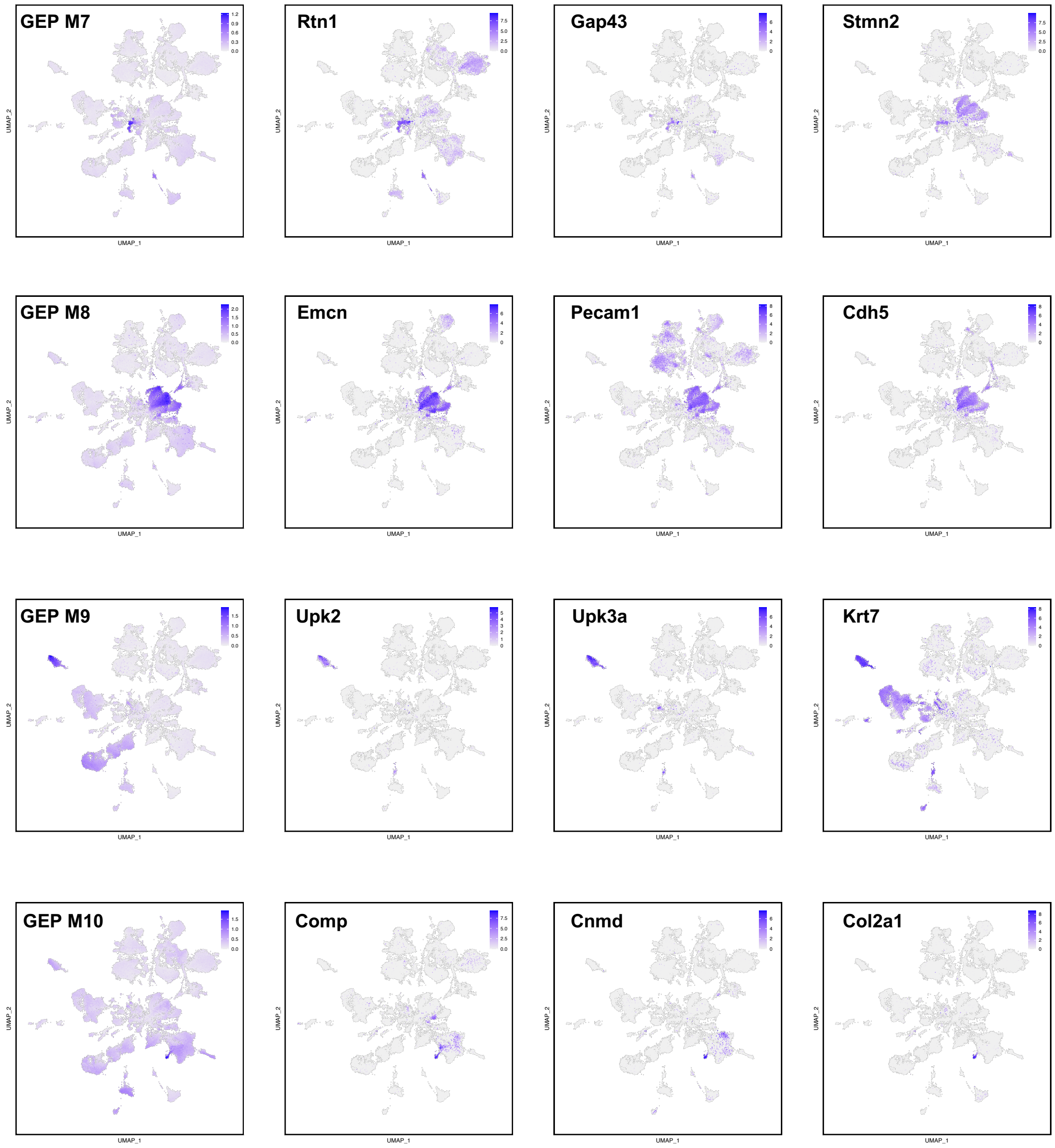

### Figure S11 continued

B

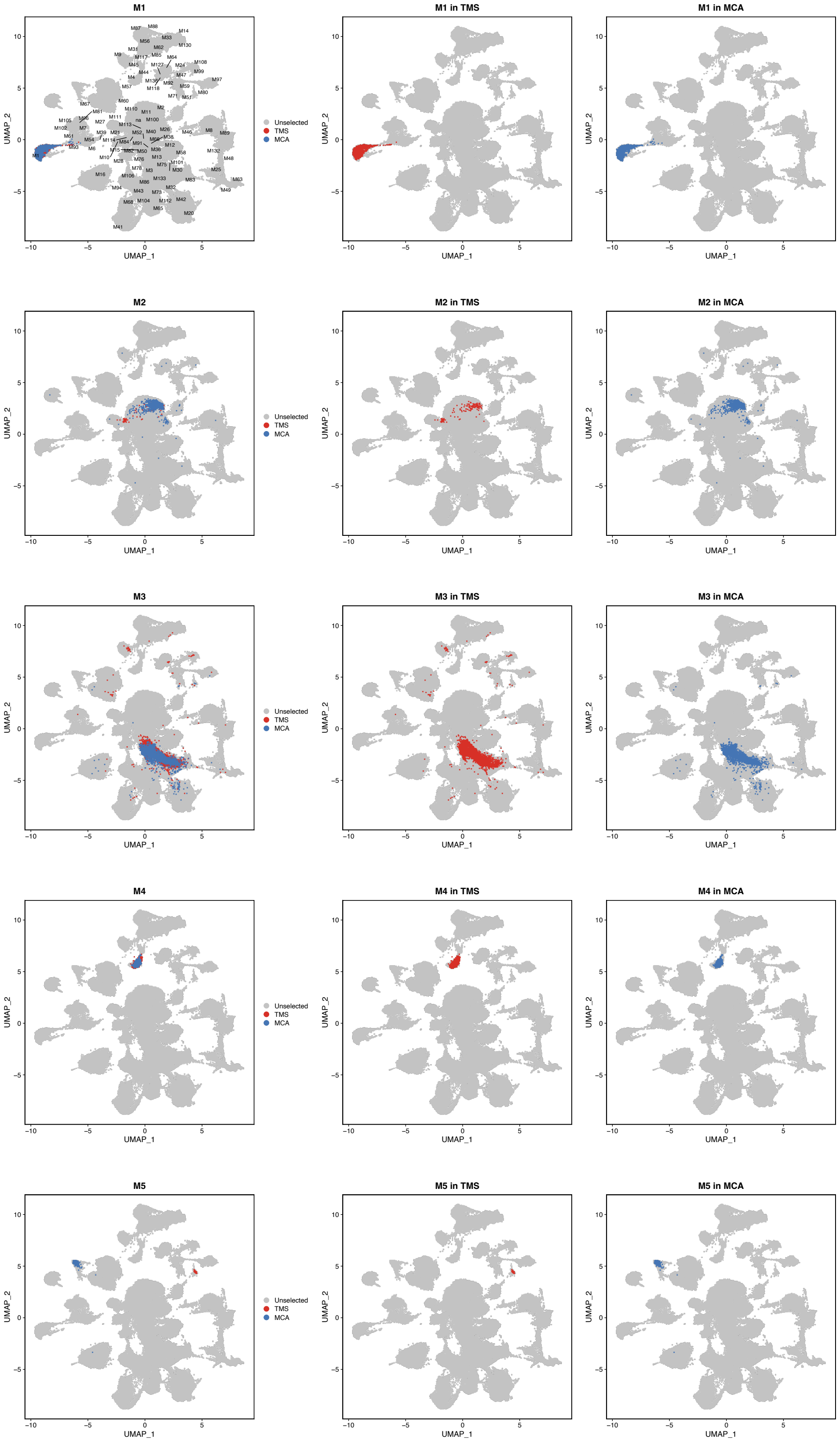

### Figure S11 continued

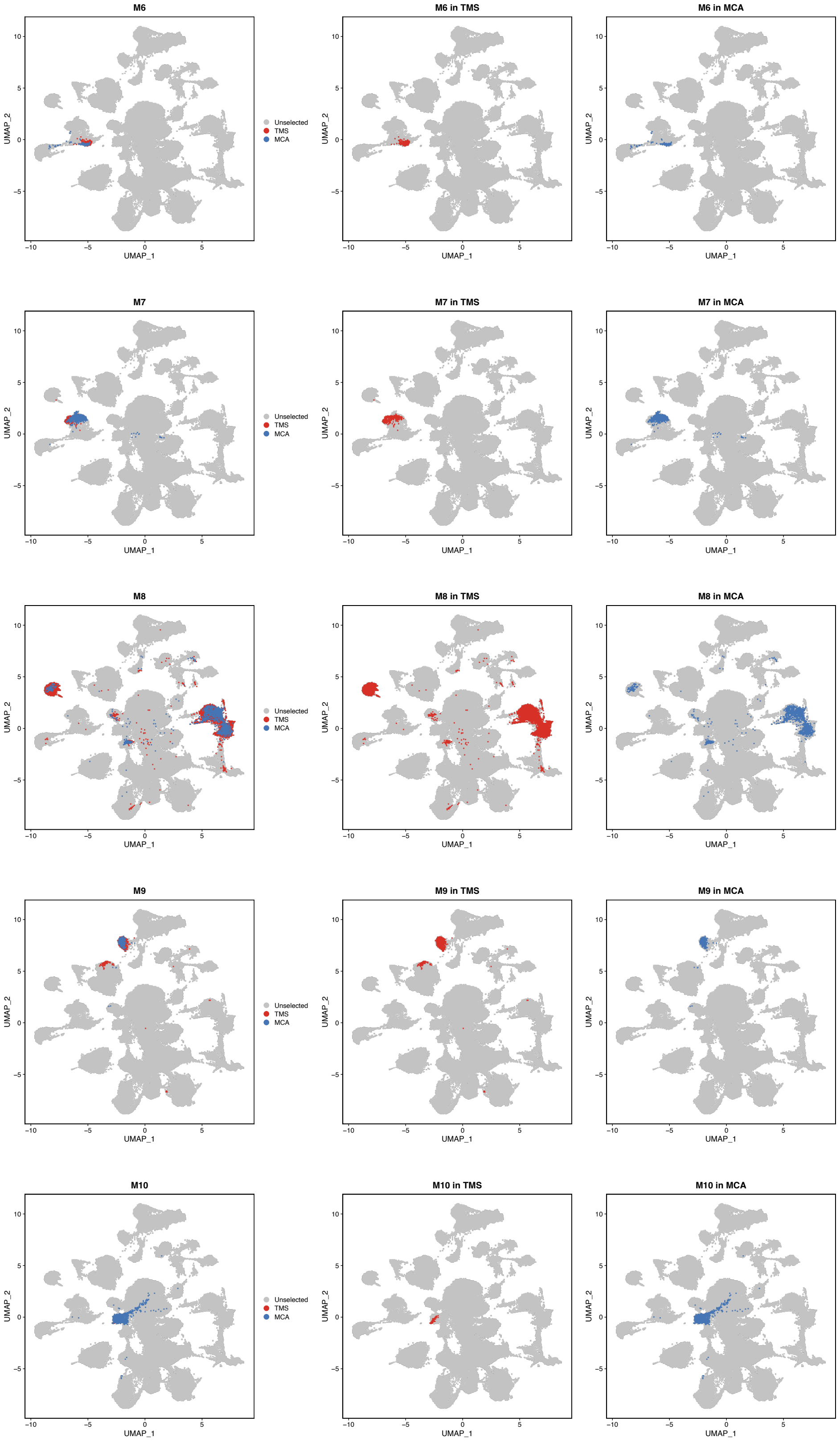

### Figure S11 continued

C

| GEP | Similar to known marker gene | Overlap in integrated UMAP |
|---|---|---|
| M1 | ✅ | ✅ |
| M2 | ✅ | ✅ |
| M3 | ✅ | ✅ |
| M4 | ✅ | ✅ |
| M5 | ✅ | |
| M6 | ✅ | ✅ |
| M7 | ✅ | ✅ |
| M8 | ✅ | ✅ |
| M9 | ✅ | ✅ |
| M10 | ✅ | ✅ |
| M11 | ✅ | ✅ |
| M12 | ✅ | ✅ |
| M14 | ✅ | ✅ |
| M16 | ✅ | ✅ |
| M17 | ✅ | ✅ |
| M20 | ✅ | ✅ |
| M21 | ✅ | ✅ |
| M23 | ✅ | |
| M24 | ✅ | ✅ |
| M25 | ✅ | ✅ |
| M28 | ✅ | ✅ |
| M30 | ✅ | ✅ |
| M31 | | ✅ |
| M32 | ✅ | ✅ |
| M33 | ✅ | ✅ |
| M40 | ✅ | |
| M41 | ✅ | ✅ |
| M42 | ✅ | ✅ |
| M43 | ✅ | ✅ |
| M44 | ✅ | ✅ |
| M48 | ✅ | ✅ |
| M49 | ✅ | ✅ |
| M51 | ✅ | ✅ |
| M54 | ✅ | ✅ |
| M56 | | |

| GEP | Similar to known marker gene | Overlap in integrated UMAP |
|---|---|---|
| M57 | ✅ | ✅ |
| M58 | ✅ | ✅ |
| M59 | ✅ | ✅ |
| M61 | ✅ | ✅ |
| M63 | ✅ | ✅ |
| M64 | ✅ | ✅ |
| M65 | ✅ | ✅ |
| M67 | ✅ | |
| M68 | ✅ | ✅ |
| M71 | ✅ | ✅ |
| M73 | ✅ | ✅ |
| M80 | | ✅ |
| M82 | ✅ | |
| M83 | ✅ | ✅ |
| M85 | | |
| M86 | ✅ | ✅ |
| M87 | ✅ | ✅ |
| M88 | ✅ | ✅ |
| M89 | ✅ | ✅ |
| M92 | ✅ | ✅ |
| M93 | ✅ | |
| M94 | ✅ | ✅ |
| M99 | ✅ | ✅ |
| M100 | ✅ | ✅ |
| M101 | ✅ | ✅ |
| M103 | ✅ | |
| M104 | ✅ | ✅ |
| M106 | ✅ | |
| M108 | ✅ | ✅ |
| M114 | ✅ | |
| M117 | ✅ | ✅ |
| M118 | ✅ | ✅ |
| M127 | ✅ | ✅ |
| M130 | ✅ | ✅ |
| M132 | | ✅ |

**Figure S11. Two lines of evidence supporting cell type label transfer between the MCA and TMS datasets. (A)** Expression of MCA GEPs and known marker genes contained in TMS cells. The left column shows the expression of marker GEPs, and the right three columns to show the expression of known marker genes contained within the GEP. Most GEP expression patterns are similar to the known marker genes. Only GEP M2 – M10 are shown due to limited space. **(B)** The cells annotated by the same GEPs from MCA and TMS after the two datasets were integrated. The cells annotated by the same GEPs from these two datasets often overlap with each other after integration. Only GEP M1-M10 are shown due to limited space. **(C)** Evidence supporting GEP identifying the same cell types in both datasets.
